# An orderly sequence of autonomic and neural events at transient arousal changes

**DOI:** 10.1101/2022.02.05.479238

**Authors:** Yameng Gu, Feng Han, Lucas E. Sainburg, Margeaux M. Schade, Orfeu M. Buxton, Jeff H. Duyn, Xiao Liu

## Abstract

Resting-state functional magnetic resonance imaging (rsfMRI) allows the study of functional brain connectivity based on spatially structured variations in neuronal activity. Proper evaluation of connectivity requires removal of non-neural contributions to the fMRI signal, in particular hemodynamic changes associated with autonomic variability. Regression analysis based on autonomic indicator signals has been used for this purpose, but may be inadequate if neuronal and autonomic activity covary. To investigate this potential co-variation, we performed rsfMRI experiments while concurrently acquiring electroencephalography and autonomic indicator signals, including heart rate, respiratory depth, and peripheral vascular tone. We identified a recurrent and systematic pattern of fMRI, EEG, and autonomic changes coincidental with intermittent arousal, suggesting arousal modulation. The temporal relationship between the various signals indicated combined neural and autonomic contribution to the fMRI signal, both of which involve widespread brain areas. The fMRI changes included brief signal reductions in salience and default-mode networks, and the thalamus, followed by a biphasic global change. These results suggest that proper measurement of functional connectivity with fMRI requires accounting for the full spectrum of autonomic and neural changes, as well as their co-variation during arousal state transitions.

## Introduction

Resting-state functional magnetic resonance imaging (rsfMRI) is widely used to chart the brain’s functional connectivity in both health and disease (Biswal et al., 1995; Fox and Raichle, 2007; Zhang and Raichle, 2010) and relies on the cerebral blood flow (CBF) response to neuronal activity (Bandettini et al., 1992; Kwong et al., 1992; Ogawa et al., 1992). While the contribution of neuronal activity to rsfMRI-derived connectivity patterns is increasingly well established (Britz et al., 2010; Brookes et al., 2011b, 2011a; He et al., 2008; Mantini et al., 2007; Musso et al., 2010), non-neurogenic CBF changes remain a potential confounding factor (Birn et al., 2009; Power et al., 2012; Shmueli et al., 2007) in the derivation of connectivity estimates.

Identifying the source contributions to the rsfMRI signal typically involves assessing temporal correlations with indicator signals. For example, a neural contribution of rsfMRI signals has been inferred by correlation with concurrently acquired electrophysiological signals (Scholvinck et al., 2010), including the alpha-band (8-12 Hz) power of electroencephalography (EEG) (Feige et al., 2005; Goldman et al., 2002; Moosmann et al., 2003). However, the rsfMRI signals also correlate with a wide variety of non-neuronal physiological measures, including respiration volume per unit time (Birn et al., 2009), peripheral vascular tone (Özbay et al., 2018), heart rate (Chang et al., 2013), arterial and venous signals (Tong et al., 2019b), cerebrospinal fluid movement (Fultz et al., 2019), and even head motion (Power et al., 2012). These correlations were often interpreted as evidence of independent, non-neurogenic contributions to rsfMRI (Das et al., 2021; Drew et al., 2020; Duyn et al., 2020; Gu et al., 2019; Keilholz et al., 2017; Power et al., 2015; Tong et al., 2019a).

Emerging evidence suggests that neurogenic and non-neurogenic contributions to the fMRI signal may not be independent, but may instead share components generated by some unitary brain process related to alertness or arousal state. For example, during light sleep, autonomic arousal was found to accompany EEG K-complex activity and fMRI signal changes (Özbay et al., 2019). Microsleep episodes detected during the resting-state fMRI scans were also associated with both EEG and physiological modulations (Soon et al., 2021). These joint neuronal and autonomic changes at arousal transitions may contribute to the rsfMRI correlations with EEG and autonomic signals, which display a similar pattern of sensory/motor dominance (Chang and Glover, 2009; Goldman et al., 2002; Shmueli et al., 2007) and also appear highly dependent on brain arousal state (Falahpour et al., 2018; Özbay et al., 2019; Yuan et al., 2013). The rsfMRI correlations with other modalities also suggest complex spatiotemporal dynamics of the underlying neural and autonomic processes. The EEG-rsfMRI correlations peak at time lags that vary across brain regions in a manner, which cannot be explained by region-specific hemodynamic delays (Feige et al., 2005; Yuan et al., 2013). This suggests that the underlying rsfMRI changes may take the form of sequential activations, as recently observed in human rsfMRI (Gu et al., 2021; Majeed et al., 2011; Mitra et al., 2015), monkey electrocorticography (ECoG) (Gu et al., 2021), and single-neuron recordings in mice (Liu et al., 2021). The associated neuronal and autonomic changes may therefore be coupled to these rsfMRI changes with systematic and distinct time delays (Fultz et al., 2019; Özbay et al., 2018; Tong et al., 2019b). Understanding these resting-state neural and autonomic dynamics could be critical to understanding their possible role in brain waste clearance (Han et al., 2021b, 2021a; van Veluw et al., 2020), to a better interpretation of the widely observed rsfMRI correlations with various modalities, and to an improved quantification of functional connectivity using rsfMRI. To approach these goals, we compared the relationship between fMRI, EEG, and autonomic indicator signals that were acquired during the resting state.

## Methods

### Experimental paradigms

Thirty-four healthy subjects were recruited in this study. Four subjects failed to complete the entire protocol and one subject was excluded because of technical issues. Two subjects were excluded because their head movement exceeded 0.5 mm mean framewise displacement (Yoo et al., 2005) over a scanning session. Thus, twenty-seven subjects (age: 22.1±3.1 years; 14 females) were included in analysis. All subjects provided informed written consent and all the procedures were approved by the Institutional Review Board at the Pennsylvania State University. Subjects were instructed to keep still and relax throughout the scans. The EEG, cardiac, respiratory, and blood oxygen level-dependent (BOLD) fMRI were simultaneously recorded during scans. Scanning sessions included a 2-min eyes-closed-eyes-open testing scan, a 5-min anatomical scan, a 10-min resting-state scan before a visual-motor task, a 15-min visual-motor adaptation task scan, and a 10-min resting-state scan after the visual-motor task.

During the eyes-closed-eyes-open scan, the subjects were directed to open and close their eyes alternatively every ∼15 seconds, repeated for five cycles. The subjects were instructed to count approximately 15 seconds, then press a button at the start of opening or closing their eyes. The eyes-closed-eyes-open scan was used to evaluate the quality of EEG data, through the presence of alpha power modulations across the two conditions. During the resting-state scans, subjects were given the option to keep their eyes either closed or open; if open, they were instructed to focus on a white fixation point (cross) at the center of a black screen. During the visual-motor adaptation task (Albouy et al., 2013), subjects manually operated a joystick to move a cursor from the center of the screen to one of eight targets located in eight radial directions. Subjects were instructed to move the cursor to arrive at the target as fast as possible. In this study, two resting-state sessions from each subject were used to investigate signals acquired during the resting state. Thus, 54 total resting-state sessions were used.

### MR data acquisition and preprocessing

MR imaging data were acquired in a 3T Prisma Siemens Fit scanner with a Siemens 20-channel receive-array coil. Foam pads were placed around the subjects’ heads to reduce motion and increase comfort. Earplugs were provided to reduce the acoustic noise during scanning. Anatomical data were acquired using a MPRAGE (magnetization-prepared rapid acquisition gradient echo) sequence with TR 2300 milliseconds, TE 2.28 milliseconds, flip angle 8 degrees, FOV 256 millimeters, 1mm isotropic spatial resolution, matrix size 256×256×192, acceleration factor 2. BOLD fMRI data were acquired using an echo planar imaging (EPI) sequence with TR 2100 milliseconds, TE 25 milliseconds, flip angle 90 degrees, slice thickness 4 millimeters, slices 35, FOV 240 millimeters, and an in-plane resolution of 3mm×3mm. Cardiac pulse data were recorded by placing a pulse oximeter (Siemens) on the left index finger with a sampling rate of 200 Hz. Respiratory effort data were recorded by placing a respiratory effort belt (Siemens) around the rib cage and abdomen with a sampling rate of 50 Hz.

We preprocessed the rsfMRI BOLD data using FCP scripts (Biswal et al., 2010) with small modifications, and the scripts used FSL (Jenkinson et al., 2012) and AFNI (Cox, 1996). The skull was removed on the anatomical image and the white-matter, gray-matter, and cerebral spinal fluid were segmented from the anatomical image. Next, the rsfMRI BOLD data were smoothed both temporally (0.01 - 0.1 Hz) and spatially (FWHM = 4 mm). Then, the anatomical image and rsfMRI BOLD data were registered to the MNI space and the nuisance parameters were regressed out, including linear and quadratic trends, motion parameters, white-matter, and CSF signals.

### EEG data acquisition and preprocessing

EEG data were acquired using a 32-channel MR-compatible EEG system (BrainAmp, Brain Products, Germany). The AFz and FCz locations were the ground and reference electrodes, respectively. The electrodes on the cap were placed according to the 10-20 International System. An electrooculography (EOG) electrode was placed under the left eye to monitor eye movement and an electrocardiography (ECG) electrode was placed on the back to record the cardiac signal. The impedances of all the electrodes were kept below 20 kΩ and the impedances of the ground and reference electrodes were kept below 10 kΩ. The raw EEG data were recorded at a sampling rate of 5000 Hz with a band-pass filter of 0 – 250 Hz.

The EEG data were preprocessed to remove the gradient artifact and ballistocardiogram artifact based on a published algorithm (Liu et al., 2012a). Briefly, the components classified as gradient artifacts from a singular value decomposition (SVD) (Golub and Reinsch, 1970) were removed for each channel. Next, the EEG data were low-pass filtered (<125 Hz) and down-sampled at 250 Hz. The pulse artifact was then removed using independent component analysis (ICA). The components from ICA were identified and removed as an artifactual component if they had a high similarity with the signal recorded from the ECG channel. The EEG channels with large residual artifacts were excluded from further analysis.

We adapted another published method (Falahpour et al., 2018) to remove data distorted by head motion. Specifically, the EEG time series from each channel were first band-filtered between 1 Hz and 15 Hz. An amplitude for each time point was derived by calculating the rms across all channels for each scanning session. Such amplitude was then down-sampled to have the same temporal resolution of rsfMRI signals. Time points with large head motion were detected if their amplitude surpassed the threshold, which was defined by summing the median number and the interquartile range multiplied by six (Devore and Farnum, 2005). A binary time series was produced by setting those detected time points to 1 and other time points to 0. The binary time series was then convolved with the hemodynamic response function from SPM (https://www.fil.ion.ucl.ac.uk/spm/) to generate a new time series, which was then binarized by using the threshold of 0. The time points detected by either one of these two binary time series were regarded as data contaminated by head motion. On average, around 3% of the time points were identified with large head motion. The subsequent analysis was conducted using the data with the time points with large head motion excluded.

### EEG spectrogram, alpha and delta power

To calculate the frequency-specific EEG power, we first calculated the spectrogram for each EEG channel using a multi-taper time-frequency transformation (window of 2 sec, step of 1 sec and tapers of 5) from Chronux (Mitra and Bokil, 2009). The mean spectrogram was calculated by averaging the spectrogram over all channels. The mean power spectrogram was then converted into decibel unit and normalized at each frequency bin by subtracting the mean and dividing by the standard deviation.

The EEG alpha modulation shown in the mean spectrogram under an eyes-closed-eyes-open session was used to confirm the data quality of EEG recorded inside the MR environment (**Fig. S1**). The mean spectrogram here was calculated by averaging only the three occipital channels since the alpha rhythm predominantly appeared at the occipital visual cortex (Berger, 1929).

To calculate alpha power, we first calculated the spectrogram for each channel and normalized by subtracting the mean and dividing by the standard deviation. Next, the normalized spectrogram was averaged within the alpha frequency band (8-12 Hz) and then was averaged across three occipital electrodes with predominantly alpha rhythm (Berger, 1929). This band-limited power signal was normalized by subtracting the mean and dividing the standard deviation, and then was down-sampled to the same temporal resolution as the rsfMRI signals, denoted as the alpha-band power. The alpha-band power was then smoothed temporally using a band-pass filter (0.01-0.1 Hz). Delta power was calculated using a similar method as alpha, but within the frequency band of 0.5-4 Hz across all channels.

### EEG-based sleep stage scoring

After removing gradient and ballistocardiogram artifacts, EEG data were re-referenced to the contralateral mastoid (or alternatively to the ipsilateral or averaged mastoids, in cases of artifact) and were bandpass filtered at 0.3-35 Hz for the identification of sleep stages. Sleep-staging of the EEG data was performed by a Registered Polysomnographic Technologist blind to other study data. Each 30-sec epoch was evaluated using data from AASM-recommended EEG exploratory derivations (F3, F4, C3, C4, O1, O2) (Iber et al., 2007) and was assigned a sleep or wake stage (W, NREM1, NREM2, NREM3, or REM).□

### Heart rate, respiratory volume and PPG amplitude

The pulse signal was first band-pass filtered (0.5-2 Hz) to increase accuracy of detecting peaks. The heart rate was calculated by averaging time differences of continuous peaks of the pulse oximetry in a sliding window of 6.3 sec centered at each rsfMRI time point and converting to units of beats-per-minute (Chang et al., 2009). The respiratory volume was calculated as the standard deviation of the respiratory trace within a sliding window of 6.3 sec centered at each rsfMRI time point (Chang et al., 2009). The respiratory volume was used in this study because it showed greater robustness to noise than the respiration volume per unit time (Chang et al., 2009). The photoplethysmography (PPG) signal was derived from the pulse oximetry. We calculated PPG amplitude by computing the root-mean-square envelop of the pulse oximetry signal and then averaging within each rsfMRI time point, similar to a previous study (Özbay et al., 2018). The heart rate, respiratory volume, and PPG amplitude were then linearly de-trended and normalized for each session by subtracting the mean and dividing by the standard deviation. The pulse oximetry signal and respiration signal were only available in 25 subjects.

### Lag-specific rsfMRI correlations with EEG alpha power and physiological signals

Voxel-wise rsfMRI correlation maps with the alpha-band power, heart rate, or respiratory volume at different time lags were calculated and averaged across all sessions. The averaged correlation maps were then converted to z-score maps with reference to control maps, which were calculated in a similar way but with temporally reversed alpha-band power, heart rate or respiratory volume. The z-scored maps were calculated by dividing the mean correlation maps by the standard error of the mean of the control maps. The function 3dFDR from AFNI (Cox, 1996) was used to correct for multiple comparisons. The cerebrospinal fluid and white-matter regions were masked out from the z-score volume map. The masks of cerebrospinal fluid or white-matter were defined based on the Harvard-Oxford subcortical structural atlas (Desikan et al., 2006) using a threshold of 70% probability. The volume maps were then converted to surface maps using Workbench (Marcus et al., 2013).

The thalamus mask was generated by taking the overlap between the thalamus region (defined from the Harvard-Oxford subcortical structural atlas) (Desikan et al., 2006) and the brain regions showing significant activation (Z > 2.5) at time lag zero in the alpha-rsfMRI correlation maps. The dorsal anterior cingulate cortex (dACC) mask was generated by taking the overlap between the anterior cingulate gyrus (defined from the Harvard-Oxford cortical structural atlas) (Desikan et al., 2006) and the brain regions showing significant activation (Z > 2.5) at time lag of zero in the alpha-rsfMRI correlation maps. The masks for sensory/motor regions include the auditory, visual or motor/somatosensory cortices, which were defined based on the Juelich histological atlas (Eickhoff et al., 2007).

### Extraction of CSF, ICA, and SSS

The CSF inflow signal, internal carotid artery (ICA) BOLD signal, and superior sagittal sinus (SSS) BOLD signal were extracted from preprocessed rsfMRI signals, including skull stripping, motion correction, and temporal filtering (0.01 - 0.1 Hz). The spatial filtering, registration to the MNI space, and regression of the nuisance parameters were skipped to focus on signals of interest and be consistent with previous studies (Fultz et al., 2019; Tong et al., 2019b). The CSF inflow signals were extracted from the CSF regions at the bottom slice of fMRI signals, which can be easily detected with much brighter signal on the T2*-weighted image, similar to a previous study (Fultz et al., 2019). The ICA and SSS regions were first roughly identified on the T1-weighted structural image using an automatic algorithm (Yao et al., 2019) and further visually inspected for the accuracy. The identified vessel masks were then registered to T2*-weighted fMRI using the FMRIB Software Library (FSL) (Jenkinson et al., 2012) to generate masks of rsfMRI data. The ICA and SSS signals were extracted from and averaged within these masks respectively. The extracted signals were further linearly de-trended and normalized for each session by subtracting the mean and dividing by the standard deviation.

### Detection of the fMRI cascade

The spatiotemporal pattern around the sensory/motor co-activation pattern was calculated as follows. For each session, we averaged rsfMRI signals within the auditory, visual, or motor/somatosensory regions respectively, and normalized each averaged time course by dividing its own standard deviation (SD). The masks for auditory, visual, and motor/somatosensory cortices were defined based on the Juelich histological atlas (Eickhoff et al., 2007). The time points with a sensory/motor co-activation pattern were defined as local peaks in the rsfMRI signal averaged from all three masks and also showing a signal amplitude larger than 1 SD in each of the three masks. (Tagliazucchi et al., 2012). A time window of 25.2 sec (12TR X 2.1 sec/TR) centered at each of selected time points with a sensory/motor co-activation pattern (set as time zero) was defined. The rsfMRI signals within these time windows were averaged to derive the spatiotemporal pattern around the sensory/motor co-activation pattern. The spatiotemporal pattern was converted to z-score maps by comparing to a null distribution, which was built by pooling rsfMRI signals within time windows centered at randomly selected time points.

To derive the fMRI cascade, we first calculated the sliding correlations between the spatiotemporal pattern around the sensory/motor co-activations and rsfMRI signals. The fMRI cascade was detected as those local peaks in the sliding correlation time course with an amplitude exceeding a pre-defined threshold. The threshold was defined as the 99.9^th^ percentile of a null distribution of sliding correlations, which were calculated between rsfMRI signals and 100 randomly selected templates from rsfMRI signals that are of equal duration as the spatiotemporal pattern. A time window of 25.2 sec (12TR × 2.1 sec/TR) centered at each of detected peaks was defined. The rsfMRI signals within these time windows were averaged to derive the fMRI cascade maps, with setting the detected peaks as time zero. The fMRI cascade maps were converted to z-score map in a similar way as the spatiotemporal pattern. The fMRI cascade maps were also computed using resting-state sessions before or after the visual-motor adaptation task, respectively (**Fig. S2**).

The temporal dynamics of various neural/physiological signals at the fMRI cascade were calculated as below. We first resampled these signals to a temporal resolution of 0.525 sec using spline interpolation to estimate accurate peak time. The various signals within the time windows centered at time points with the fMRI cascade were averaged to derive their temporal dynamics at the fMRI cascade. The global BOLD (gBOLD) signals were calculated by averaging the rsfMRI signals within the cortical gray-matter region using rsfMRI signals without regressing out the CSF, global signal, or white-matter signal. The masks of cortical gray-matter, or white-matter were defined based on the Harvard-Oxford subcortical structural atlas (Desikan et al., 2006) using a threshold of 70% probability.

The dependency of the fMRI cascade on brain states was examined by categorizing and averaging signals within various brain states, which were defined based on EEG signals mentioned above. Specifically, each time window around the time point with the fMRI cascade was categorized as one of the sleep stages or awake states if the brain state within the time window was assigned a single brain state.

### Removing the dynamics of fMRI cascade

The temporal dynamics of the fMRI cascade was regressed out from the rsfMRI signals using a published algorithm (Abbas et al., 2019). Briefly, an fMRI cascade time course was calculated for each voxel by convolving the fMRI cascade dynamics, i.e., the sliding correlations between the fMRI cascade and rsfMRI signals, with this voxel’s temporal dynamics during the fMRI cascade. We then used a linear regression model to remove the temporal dynamics of the fMRI cascades by regressing out the voxel-specific fMRI cascade predictor. The effect of fMRI cascade was also regressed out from other signals in a similar way, including the alpha power, HR, RV, PPG amplitude, CSF, ICA, and SSS. The correlations between different signals were then calculated using the signals with the fMRI cascade dynamics removed. It should be noted that the rsfMRI signals used to calculate their correlations with the PPG amplitude did not regress out the white-matter signal in order to keep the signal of interest since the rsfMRI signals in the white-matter showed systematic correlation with the PPG amplitude (Özbay et al., 2018).

## Results

### Similar rsfMRI correlations with EEG alpha power and physiological signals

We analyzed concurrently acquired fMRI, EEG, and autonomic signals from 27 subjects (14 females) at rest in the scanner. First, we studied the fMRI correlations with EEG alpha activity, a correlate of alertness/arousal (Makeig and Inlow, 1993; Putilov and Donskaya, 2014). Similar to previous studies, positive correlation was seen in the dorsal anterior cingulate cortex (dACC) and thalamus (de Munck et al., 2007; Feige et al., 2005; Liu et al., 2012), as well as in the insula, that appeared to peak at lags much shorter than the neurovascular delay (**Fig. 1A**). These brain regions are often considered major components of the salience network (SN; 91% overlapped with Neurosynth-defined SN) (Seeley et al., 2007) (**Fig. S3, A-B**). The dACC reached its peak positive correlation with EEG alpha-power at a slightly shorter time lag (∼0 sec) as compared with the thalamus (∼2.1 sec) (**Fig. 1B**). Shorter-lag positive correlations were also seen at the medial frontal gyrus (MFG) and posterior cingulate cortex (PCC), which are areas significantly overlapping with the default-mode network (DMN; 88% overlapped with Neurosynth-defined DMN) (Raichle, 2015; Raichle et al., 2001). At lags slightly longer than the neurovascular delay (at ∼7-sec as compared with the canonical “neurovascular delay” of 5-6s (Buxton et al., 2004)), widespread negative correlations were observed, most strongly in sensory/motor regions, including visual, motor, auditory, and somatosensory cortices (**Fig. 1A**). The different lag dependence of the correlations is clearly visible across brain regions as shown in **Fig. 1B**. In summary, resting-state EEG alpha-power changes were significantly associated with rsfMRI changes at brain-region-dependent time lags, with the earliest effects (shortest lag correlations) seen at the SN and DMN.

**Fig. 1.**
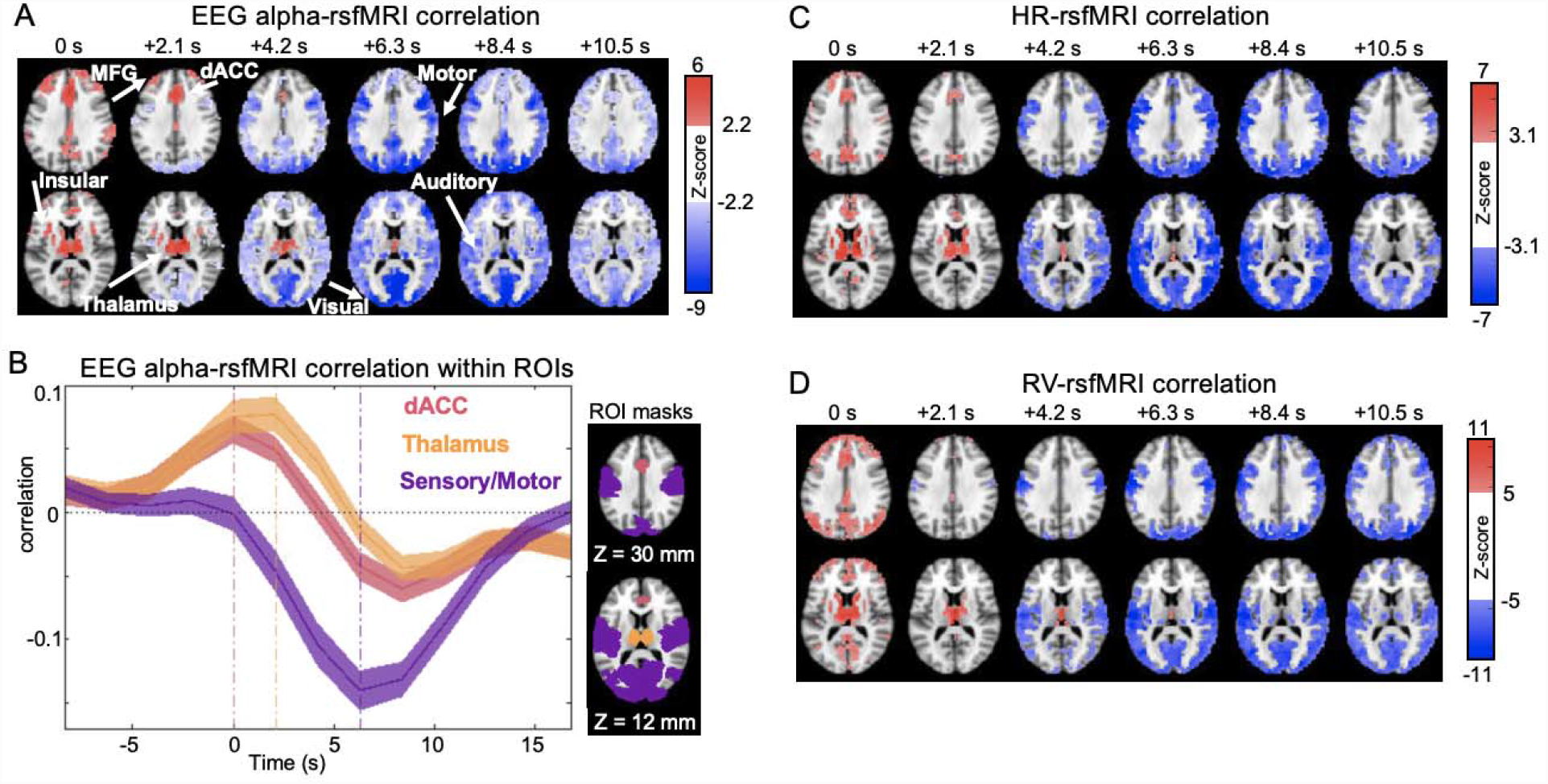
Lag-dependent rsfMRI correlations with the EEG alpha power and physiological signals showed similar patterns. (A) The correlations between rsfMRI and EEG alpha-power (8-12 Hz) show distinct spatial patterns at various time lags. Average correlations across 54 sessions were converted to z-scores and thresholded at an FDR-corrected q value of 0.05. The rsfMRI was used as the reference, and thus the positive lags correspond to delayed EEG alpha-power. (B) The lag-dependent rsfMRI correlations with the EEG alpha power were averaged within three regions of interest (ROIs) across 54 sessions. The shaded regions represent the area within 1 S.E.M. (C) The rsfMRI correlations with the heart rate (HR), which were thresholded at an FDR-corrected q value of 0.005. (D) The rsfMRI correlations with the respiratory volume (RV), which were thresholded at an FDR-corrected q value of 5×10^−7^.

Next, we examined rsfMRI correlations with autonomic indicators, specifically the instantaneous heart rate (HR) and respiratory volume (RV). Previously, these autonomic indicators were found to correlate with fMRI, and were attributed to autoregulatory control of cerebral blood flow (Birn et al., 2009; Shmueli et al., 2007). Both HR-rsfMRI and RV-rsfMRI correlations showed lag-dependent patterns highly similar to the EEG-rsfMRI correlations, with spatial correlation values peaking at 0.80 and 0.71, respectively. With both HR and RV, significant positive correlations were observed in the SN and DMN at a time lag of 0 sec, whereas negative correlations in the sensory/motor regions peaked at ∼7 sec (**Fig. 1, C-D** and **Fig. S3, C-D**). Overall, these correlation patterns were quite similar to those seen with the EEG alpha-power correlation (**Fig. 1A**), suggesting a strong coordination of changes across the various signals through a unitary phenomenon, apparently related to arousal transitions indicated by EEG alpha-power modulations.

### A cascade of fMRI dynamics contributes to rsfMRI correlations with EEG and autonomic physiology

The lag-dependent rsfMRI correlations with EEG and cardio-respiratory physiology suggest that underlying fMRI activity may involve sequential modulations of the SN/DMN and sensory/motor regions. However, it is also possible that the two sets of brain regions were linked to EEG and HR and RV changes through independent brain processes, and thus changed independently from each other. To investigate this, we aligned and averaged rsfMRI time segments with respect to time points showing a strong sensory/motor co-activation pattern (see the Methods for detail) that resembled the EEG-rsfMRI correlations at the 6.3-sec time lag (**Fig. 1A)**. The SN and DMN were found to de-activate ∼7.9 and ∼7.4 sec, respectively, prior to the sensory/motor co-activations (**Fig. S4**), forming a cascade of fMRI dynamics with sequential modulations in the SN/DMN and sensory/motor regions. We used this spatiotemporal pattern as an initial template to detect and refine the fMRI cascades (**Fig. 2A** and **Fig. S3, E**) (see the Methods for detail), resulting in the final cascade pattern that was highly similar (*r* = 0.98) to the original template (**Fig. S4**). Over the course of the fMRI cascade, the EEG alpha power, HR, and RV all showed strong co-modulations (**Fig. 2B**), consistent with the strong temporal correlations observed between fMRI and all other signals (**Fig. 2, C-E**). More importantly, regressing out the temporal dynamics of the fMRI cascade reduced these correlations to a large extent (by 93.9% for EEG alpha-rsfMRI peak covariance, 99.5% for HR-rsfMRI peak covariance, and 99.6% for RV-rsfMRI peak covariance) (**Fig. 2, C-E**), suggesting that the fMRI cascade led by the SN/DMN changes was their main contributor.

**Fig. 2.**
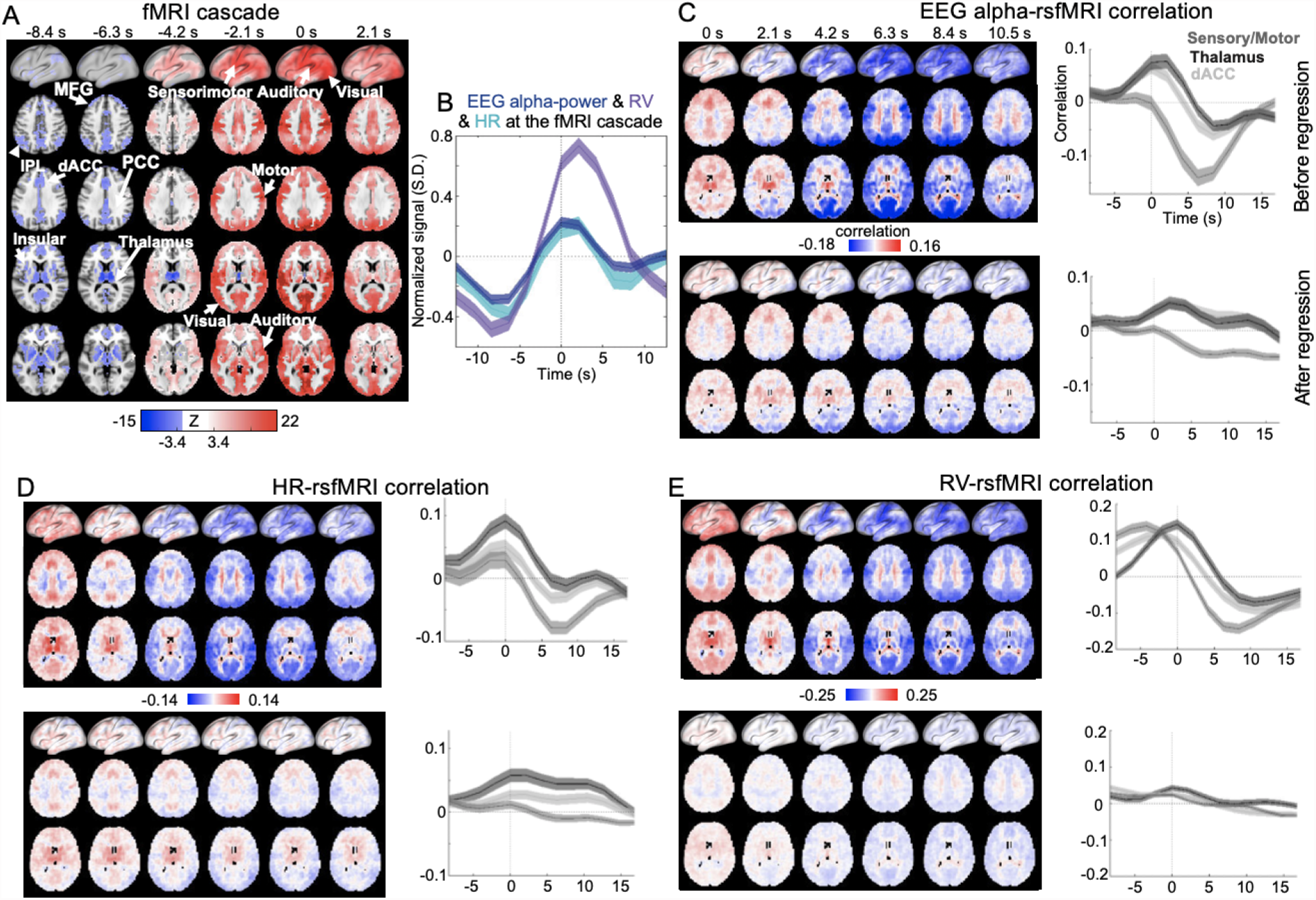
Reduction of rsfMRI correlations with EEG and physiology signals after regressing out the fMRI cascade dynamics. (A) The fMRI cascade computed by averaging fMRI time segments (N=338) with a significant (p < 0.001, permutation test) correlation with the spatiotemporal pattern with a sensory/motor co-activation pattern in Fig. S4. The fMRI cascade was converted to z-scored map and thresholded at an FDR-corrected q value of 5×10^−4^. (B) The EEG alpha-power, HR, and RV changes at the fMRI cascade. (C) The averaged alpha-rsfMRI correlation maps (left), as well as the cross-correlation functions in the three representative regions (right), before (top) and after (bottom) regressing out the fMRI cascade. (D-E): The HR/RV-rsfMRI correlation maps and cross-correlation functions before (top) and after (bottom) regressing out the fMRI cascade. IPL, inferior parietal lobe.

### EEG changes around the fMRI cascade

We previously described a prototypical time-frequency pattern of intracranial EEG changes around peaks in the fMRI global signal in the form of sequential spectral transitions (SSTs) (Liu et al., 2018, 2015) interpreted as arousal modulations. To investigate the relationship of SSTs with the fMRI cascade described here, we averaged EEG spectrogram segments time-aligned to the cascade. Similar to SSTs, the EEG activity associated with fMRI cascades showed an initial large alpha-beta (7-25 Hz) power reduction that was followed by a bout of delta-band (0.5-4 Hz) power increase (**Fig. 3A**). This pattern was then followed by a rebound of alpha-beta power (**Fig. 3A**), suggesting a rebound of arousal after the initial drop.

**Fig. 3.**
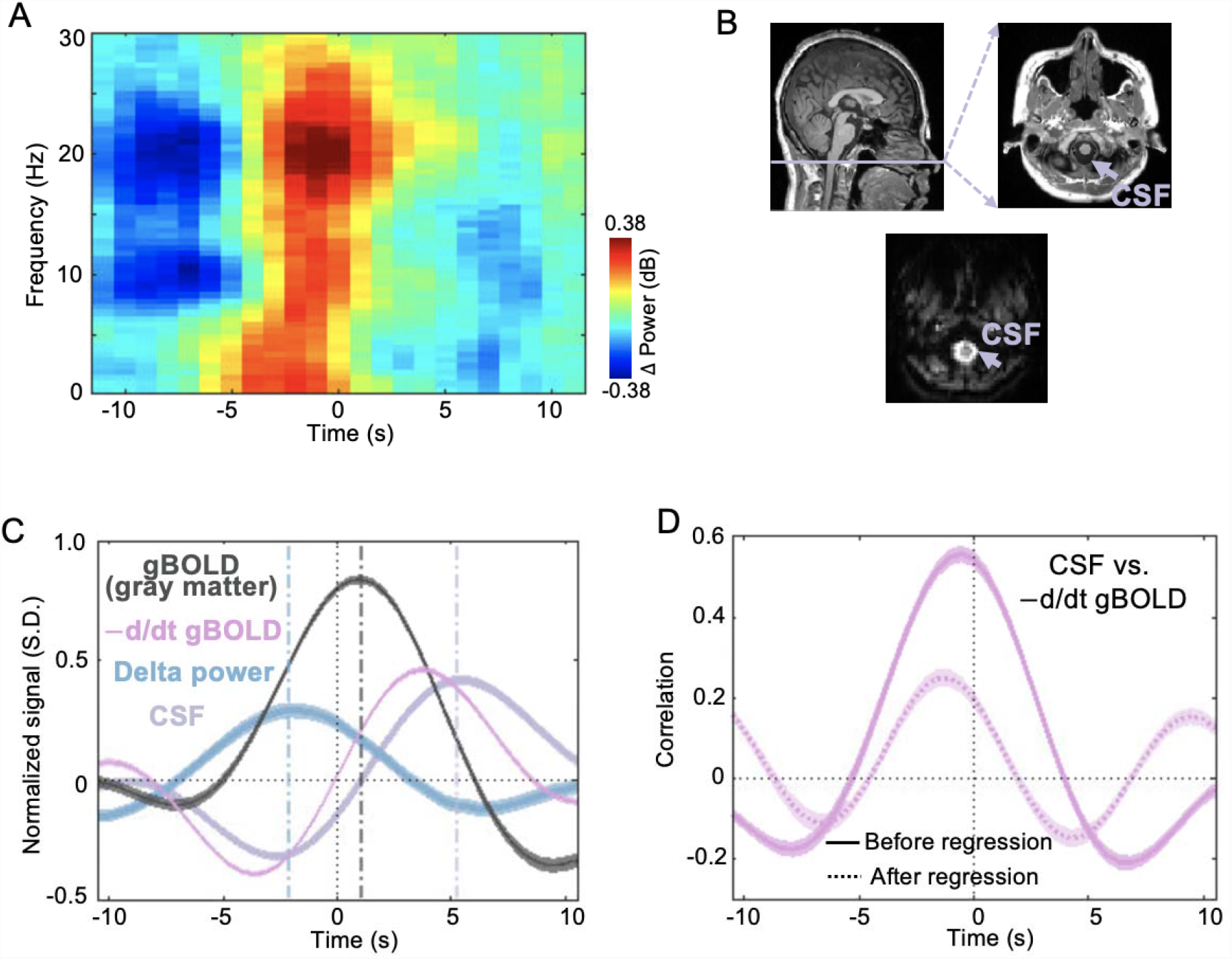
Changes in frequency-specific EEG power and CSF at the fMRI cascade. (A) The averaged EEG time-frequency modulations at the fMRI cascade. It was obtained by aligning and averaging EEG spectrogram at fMRI cascade segments. (B) The CSF region was identified as the bright voxels on the bottom slice of functional image, which corresponded to the CSF region in the T1-weighted image. (C) The global gray-matter BOLD (gBOLD), the negative derivative of gBOLD signal, EEG delta power, and CSF inflow signal changes at the fMRI cascade. (D) The averaged cross-correlation of the negative derivative of gBOLD signal with the CSF inflow signal across 54 sessions before and after regressing out the fMRI cascade dynamics. The negative derivative of gBOLD signal was used as the reference signal. The shaded regions represent area within 1 S.E.M.

### Global fMRI and CSF fluctuations around the fMRI cascade

The global BOLD (gBOLD) and the EEG delta-band power, which both co-occur with the SST (Liu et al., 2018, 2015), were recently found coupled to the cerebrospinal fluid (CSF) flow (Fultz et al., 2019). This gBOLD-CSF coupling was further linked to Alzheimer’s disease pathology and cognitive decline in Parkinson’s patients, presumably due to its role in glymphatic clearance (Han et al., 2021b, 2021a). We therefore investigated the potential link of the fMRI cascade to the global signal and CSF pulsations. The CSF flow was measured from the bottom slice of fMRI acquisition volume, analogous to previous work (Fultz et al., 2019). The results show the gBOLD signal averaged within the grey-matter region peaking at around 1s, and a bipolar CSF signal centered on this signal (**Fig. 3C**). In addition, the CSF flow signal closely followed the negative derivative of the gBOLD signal (**Fig. 3C**), an indicator of CBV change, as expected from the assumption of constant total brain fluid volume (Fultz et al., 2019). Consistent with this, a strong correlation was observed between these two signals, which was dramatically reduced (82.9% reduction in the peak co-variance) after regressing out the cascade dynamics (**Fig. 3D**). This suggests a substantial contribution of the cascade phenomenon to both the fMRI global signal and CSF pulsations.

### Vessel signals around the fMRI cascade

RsfMRI signals are also linked to other peripheral physiology measures, including the photoplethysmography (PPG) amplitude of cardiac signals (Özbay et al., 2018) and near-infrared spectroscopy (NIRS) measure of hemoglobin concentration at fingertips (Tong et al., 2012). The associated rsfMRI changes demonstrate systematic time delays across large arteries and veins (Tong et al., 2019b), as well as between the gray and white matters (Özbay et al., 2018). We thus examined the temporal dynamics of the PPG amplitude and fMRI vessel signals over the course of the fMRI cascade. The PPG amplitude had a big drop at the late phase of the fMRI cascade (4.7 sec) that was much more delayed than the largest HR modulations (the negative peak at –6.8 sec) (**Fig. 4A**). The gray-matter and white-matter rsfMRI signals showed very similar temporal dynamic but with a significant delay of ∼1.5 sec (**Fig. 4A**). Their cross-correlation functions with the PPG amplitude also showed patterns (**Fig. 4B-4C, Fig. S5**) similar to the previous report (Özbay et al., 2018). These correlations however decreased significantly (by 83.5% and 93.0% for the gray-matter and white-matter fMRI correlations respectively) after regressing out the fMRI cascade dynamics (**Fig. 4B-4C, Fig. S5**).

**Fig. 4.**
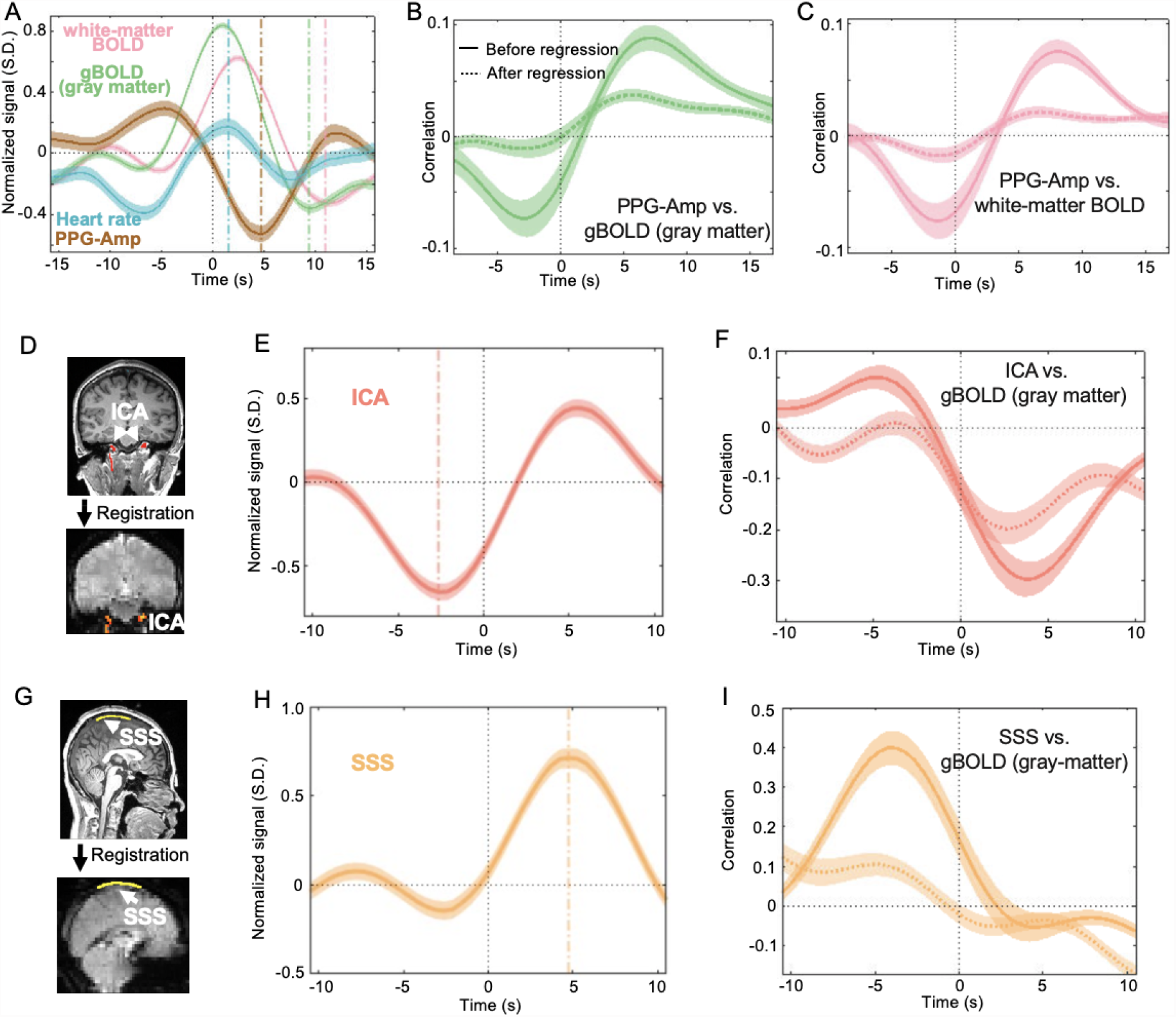
The dynamics of peripheral PPG and vessel signals at the fMRI cascade. (A) The modulations of heart rate, PPG amplitude, rsfMRI signals within gray-matter and white-matter at the fMRI cascade. (B-C) The averaged cross-correlation of PPG amplitude with the rsfMRI signals within gray-matter and white-matter across 54 sessions before and after regressing out the fMRI cascade dynamics. The gray-matter or white-matter rsfMRI signals were used as the reference signal. (D, G) The ICA and SSS regions were directly identified on the T1-weighted image and then registered to the functional image to extract vessel signals. (E, H) The modulations of ICA and SSS signals at the fMRI cascade. (F, I) The averaged cross-correlation of the global gray-matter BOLD (gBOLD) signals with ICA or SSS signals across 54 sessions before and after regressing out fMRI cascade dynamics. The gBOLD signal was used as the reference signal. The shaded regions represent area within 1 S.E.M.

The rsfMRI signals of large blood vessels, including internal carotid artery (ICA) and the superior sagittal sinus (SSS) (**Fig. 4D** and **4G**), showed strong but distinct modulations across the fMRI cascade. The ICA signal showed a big drop at –2.6 seconds whereas the SSS signal displayed a large positive peak at 4.7 seconds (**Fig. 4E** and **4H**). The ICA and SSS signals were significantly correlated with the gBOLD signal, with a distinct peak offset consistent with these relative delays and also with previous reports (Tong et al., 2019b). Such gBOLD-vessel correlations reduced significantly (by 59.5% and 93.6% for the ICA and SSS respectively) after accounting for the temporal dynamics of the fMRI cascade (**Fig. 4F** and **4I**), suggesting that they were largely caused by their co-modulation during the fMRI cascade.

### Dependency of the fMRI cascade on brain state

To investigate the dependency of fMRI cascade results on sleep/wake brain states, we split and examined the data according to sleep stages scored from the EEG signals. Among all the identified fMRI cascade events, 43% occurred during the wakefulness whereas 20% and 17% were found at the sleep stages 1 and 2, respectively. The remaining events happened during the time periods with mixed wake and sleep (stages 1 and 2 only). The fMRI cascades showed similar overall patterns across wake and the different sleep stages, except for the SN/DMN de-activations (**Fig. 5**). The SN/DMN de-activation at the early phase of the fMRI cascade showed a clear dependency on the brain state, with a gradual reduction from awake to sleep stage 1 and then to sleep stage 2 (**Fig. 5A**). In contrast, the de-activation of the thalamus, which occurred at the similar phase as the SN/DMN de-activations, remained similar across different states. The EEG signals showed large changes in the delta-band activity in sleep stage 2 as compared with the other two states (**Fig. 5B**). The gray-matter gBOLD signal, white-matter BOLD signal, ICA signal, and PPG-Amp showed similar amplitude of changes across wakefulness and different sleep stages, whereas the HR, RV, CSF flow signal, and SSS appeared to display larger changes during sleep as compared to wake (**Fig. 5C**).

**Fig. 5.**
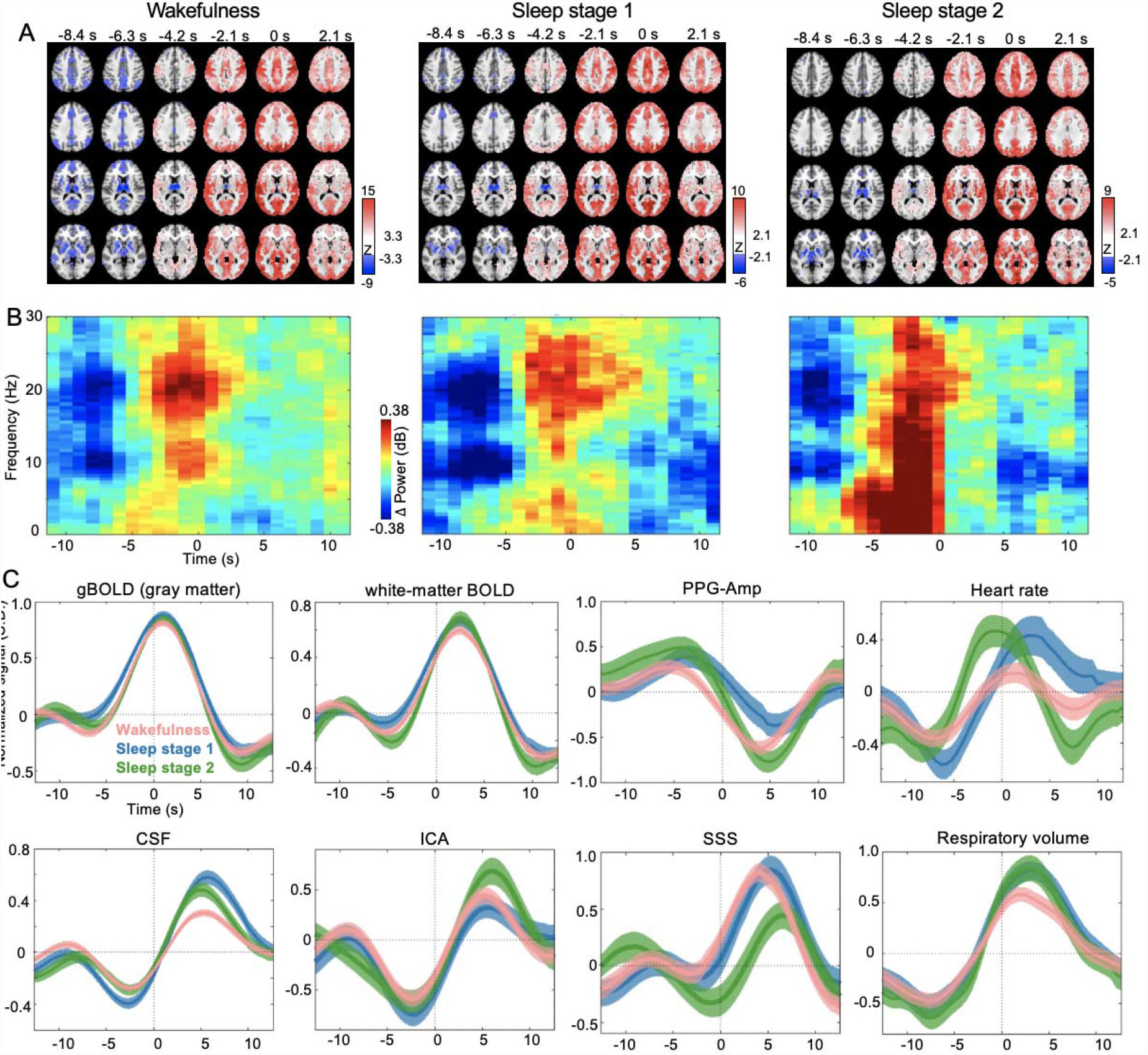
The various signals at the fMRI cascade were categorized by wake and different sleep stages. (A) The fMRI cascade was grouped and averaged within various brain states: the wake (43% of the total), sleep stage 1 (20%) and sleep stage 2 (17%). (B-C) The EEG time-frequency modulations and temporal dynamic of other physiological signals at the fMRI cascade were also grouped and averaged within various brain states.

### Temporal order of the multimodal modulations at the fMRI cascade

The availability of multimodal data allowed us to summarize the temporal order of neural and physiological modulations at the fMRI cascade (**Fig. 6**). Assuming a neurovascular delay of 5.6 sec (Buxton et al., 2004), putative neuronal changes in SN and DMN showed the earliest modulations at –13.5 sec and –13 sec respectively with respect to the fMRI cascade center (same hereinafter), which were followed by the reductions in the respiratory volume (–7.4 sec), the heart rate (–6.8 sec), and the EEG alpha-power (–6.8 sec) around –7 sec. The putative neuronal activity underlying the gBOLD peak (1.1 sec) would occur around –4.5 sec, which is slightly delayed with respect to the EEG alpha-power reduction but is earlier than the EEG delta-power increase at –2.1 sec. The fMRI changes followed a specific sequence, from earliest changes in the ICA (–2.6 sec), followed by gray matter (1.1 sec), white matter (2.6 sec), and eventually to SSS (4.7 sec), though the ICA signal showed opposite changes to others. Finally, the PPG amplitude reduction (4.7 sec) and the CSF signal changes (5.3 sec) occurred at the relative late phase of the fMRI cascade.

**Fig. 6.**
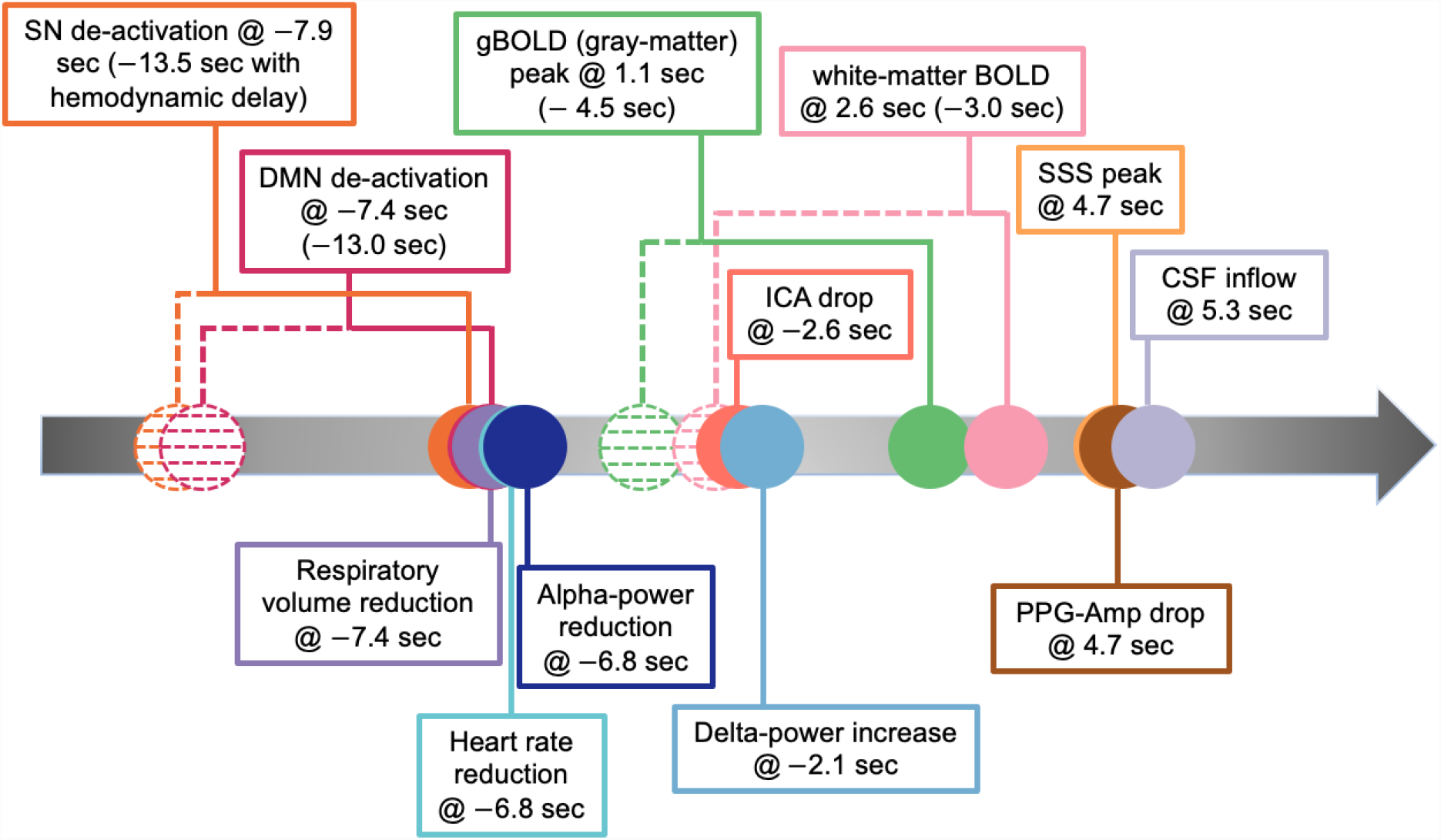
The temporal order of the multimodal modulations at the fMRI cascade.

## Discussion

In experiments performed on resting human subjects, we observed joint fluctuations in fMRI, electro-cortical and autonomic activity. A prototypical spatio-temporal pattern of fMRI changes was observed to coincide with a biphasic modulation of EEG band-limited power in the alpha/beta range as well as with biphasic changes in autonomic indicator signals.

The nature of the signal changes indicated that they originate from an initial drop in cortical arousal, closely followed by a rebound. Early fMRI changes in regions often associated with arousal modulation, followed by reduction in mid-frequency EEG band-limited power, and reduced heart rate and respiratory depth all suggest a drop in arousal level. These early neurogenic changes are consistent with the role of the salience network in maintaining tonic attention (Sadaghiani and D’Esposito, 2015). In fact, these de-activated regions are largely overlapped with the brain areas showing significantly reduced glucose metabolism and CBF at the dexmedetomidine-induced unconsciousness (Akeju et al., 2014). Interestingly, the SN/DMN deactivations were much weaker in the fMRI cascades obtained during sleep, which might be due to the already low activity level in these cortical regions during sleep. The subsequent widespread increase in fMRI signal, mid-frequency EEG power reduction, and autonomic activity decrease are suggestive of an arousal drop. The late, widespread decrease in fMRI signal is consistent with previously reported effects of increased autonomic activity that are associated with the arousal rebound, including increases in respiratory depth (Birn et al., 2008a) and sympathetic vasoconstriction (Özbay et al., 2018).

A possible mechanistic explanation for the joint occurrence of changes in EEG, fMRI, and autonomic indicators is the well-established close interaction between the ascending reticular activation system and the autonomic regulatory centers in the brain stem (de Zambotti et al., 2016; Duyn et al., 2020; Silvani et al., 2015). This close interaction is not surprising, given that cortical and autonomic arousal are mediated from overlapping neural substrates. Joint activation may arise from internal or external stimuli through mid-brain relays, or initiated by a cognitive process (Dampney, 2015). This mechanism may explain previous observation of a strong dependency of EEG-rsfMRI and physiology-rsfMRI correlations on brain state (Falahpour et al., 2018; Yuan et al., 2013), as well as the joint EEG/fMRI/autonomic changes that have also been observed during light sleep (Özbay et al., 2019).

These findings significantly impact the interpretation of previous fMRI experiments. They suggest that signal correlations may relate to a unitary brain process that affects both electro-cortical and autonomic activity, making it difficult to assign the relative contribution of neuronal and autonomic sources to the fMRI signal. In addition, both contributions appear to be widespread, complicating the interpretation of interregional signal correlations in terms of their functional connectivity. Lastly, the observed joint changes in autonomic and neuronal activity are likely not specific to rsfMRI, and might be expected to occur in task-based fMRI as well. In fact, previous research indicated the prevalence of task-induced autonomic changes in a variety of cognitive and motor tasks (Glasser et al., 2018).

The cascade of fMRI signal changes observed here and attributed to an arousal transition may similarly have contributed to previous findings. For example, the fMRI cascade clearly overlaps with the gray-matter BOLD peaks given its similar sensory-dominant pattern and the associated EEG changes of an SST pattern (**Fig. 3A**), which has been observed in monkey electrophysiology at the large gBOLD peaks previously (Liu et al., 2018). In fact, the SST has previously been linked to transient (∼10 seconds) modulations of brain arousal state (Liu et al., 2018, 2015). Previous rsfMRI studies of spontaneous eye closure also found multiphasic signal behavior, with changes in the thalamus appearing to differ (Chang et al., 2016) and precede (Soon et al., 2021) most of cortical grey matter. Arousal transitions and their associated fMRI signal cascade may also have contributed in previous reports of spatiotemporal rsfMRI structures, such as quasi-periodic patterns (QPP) (Majeed et al., 2011) and cross-hierarchy propagations (Gu et al., 2021), both of which showed sequential fMRI co-(de)activations at different brain regions.

The fMRI cascade, at least in part, appears to result from cortical neuronal activity. First, it shows a specific sequence of network involvements, i.e., the sensory/motor co-activations preceded by the SN/DMN deactivations. It seems unlikely that the non-neuronal physiological changes would lead to such a highly organized pattern of networks. Second, the cascade is associated with strong EEG changes featured by sequential modulation of the alpha-beta and delta powers. In fact, this time-frequency pattern is similar to the SST event observed with the gBOLD peaks (Liu et al., 2018). At the single-neuron level, this fMRI cascade could be linked to a temporal cascade of spontaneous spiking activity of large neuronal populations recently found in mice (Liu et al., 2021). This spiking cascade occurring during the resting state is of a similar time scale (5-20 seconds), leading to big peaks in the global mean spiking rate, and accompanied by strong delta-power modulations and sympathetic responses measured as pupil dilation. In fact, the spiking cascade takes the form of sequential activations between two distinct neuronal groups that showed opposite modulation across the running and resting states. It is possible that the fMRI cascade at the network level might result from an uneven distribution of the two types of neurons at the SN/DMN and sensory/motor networks.

In addition to neuro-vascular source, it appears that autonomic mechanisms also contribute to the observed fMRI signal changes. For example, the timing of the late, widespread reductions in fMRI signal at the end of the cascade (**Fig. S6**) relative to that of the autonomic arousal (evidenced from the joint drop in PPG-amplitude and increase in HR and RV) are consistent with the sympathetic mechanism described previously (Özbay et al., 2019). A similar autonomic-fMRI relationship was observed in a recent study of microsleeps defined by periodic eye closure (Soon et al., 2021). Judging from the PPV-amplitude changes, as well as the relatively short (∼10s) delay between RV and fMRI changes that was consistent with the peak delay of respiratory response function (RRF) during resting state but shorter than the ∼16s delay of the RRF induced by cued deep breathing (Birn et al., 2008b; Chang et al., 2009), sympathetic vasoconstriction may be the main contributor to the late, widespread signal reductions in the fMRI cascade.

Less clear is the origin of the gBOLD peak (at 1.1s in the cascade). This peak may not be directly induced by the EEG alpha-power reduction given the fact their relatively delay (∼7.9 sec) is slight longer than the neurovascular delay. One possibility is that it is linked to increases in gamma-power (>40 Hz) spectral frequencies that are poorly visible in scalp EEG here but evident in ECoG-measured SSTs (Liu et al., 2018). Like the SST gamma-power increase, the gBOLD peak in the cascade shows a sensory-dominant co-activation pattern (Liu et al., 2018, 2015), and its ∼5.8s delay after a putative gamma power increase (which was estimated as 2.1 sec after the alpha-beta power reduction in the SST (Liu et al., 2015)) would be more consistent with the canonical hemodynamic delay.

The relatively strong (and rapid) autonomic rebound accompanying the apparent rebound in cortical arousal may also have a homeostatic role. Previous work has suggested a relationship between respiratory and cardiac cycles and CSF pulsations that support brain waste clearance (Iliff et al., 2013a; Klose et al., 2000; Stoodley et al., 1997; Yamada et al., 2013). Physiological changes, particularly CSF movement, could be critical for a glymphatic system that relies upon CSF flow through the perivascular and interstitial spaces to clear brain waste, such as amyloid-beta (Iliff et al., 2012; Tarasoff-Conway et al., 2015). Though arterial pulsations have traditionally been regarded as the major driving force for the glymphatic CSF flow (Iliff et al., 2013b; Schley et al., 2006), these pulsations are mediated by changes in cerebral vascular tone secondary to changes in arterio-venous blood pressure differences and are often weak during sleep (Baust and Bohnert, 1969; Boudreau et al., 2013; Douglas et al., 1982; Snyder et al., 1964), which is in discordance with the sleep-enhanced glymphatic clearance (Xie et al., 2013). Recent reports suggest that an important contribution to CSF pulsations may originate from slow (<0.1 Hz) changes in cerebral vascular tone (Fultz et al., 2019), with possible contribution from neurovascular (van Veluw et al., 2020) or autonomic mechanisms (Özbay et al., 2018; Picchioni et al., 2021). In addition, the coupling between such CSF pulsations and gBOLD, which was shown to be strongly related to the fMRI cascade in this study, has been found associated with Alzheimer’s disease (AD) pathology (Han et al., 2021b) and Parkinson’s disease (PD) cognitive decline (Han et al., 2021a). The neuronal modulation at the fMRI cascades might coordinate with the associated autonomic changes and further facilitate the glymphatic clearance.

Regressing out the cascade-related signal resulted in strong reduction in the HR-fMRI and RV-fMRI correlations (**Fig. 2**). This suggests that previously observed HR-fMRI (Shmueli et al., 2007) and RV-fMRI (Birn et al., 2006) correlations may, in part, have resulted from intermittent changes in arousal state. Therefore, care has to be taken with removing the effects of HR and RV from the fMRI signal based on regression analysis (Birn et al., 2008b; Chang et al., 2009), because this may also result in accidental removal of the neurogenic contributions to the fMRI signal related to cortical arousal changes. Careful accounting for spatio-temporal differences between neurovascular and autonomic contributions to the fMRI signal with arousal changes may help distinguish them, and appears critical in interpreting resting state fMRI experiments.

## Author Contributions

Conceptualization, Y.G. and X.L.; Data Collection, Y.G., F.H., L.E.S., and X.L.; Data Preprocessing, Y.G., F.H., M.M.S., and X.L.; Methodology, Y.G. and X.L.; Formal Analysis, Y.G., M.M.S., and X.L.; Original Draft: Y.G., J.H.D., and X.L.; Review & Editing, Y.G., M.M.S., O.M.B., J.H.D., and X.L.; Visualization, Y.G., and X.L.; Supervision, X.L.; Funding Acquisition, X.L.

## Acknowledgments

We would like to express special thanks to Dr. Thomas Beck and Dr. Josef Pfeuffer, Siemens AG, Healthcare Sector for providing us with the Advanced fMRI package WIP package. We would also like to thank Dr. Xiaoxiao Bai, Jack W. Williams and Youyou Yu for their help during the experiment. Data analysis were conducted using the computing resources provided by the Institute for Computational and Data Sciences at the Pennsylvania State University (https://icds.psu.edu).

## Data availability

The EEG and fMRI data are available on the OpenNeuro website and can be assessed through the dataset DOI: 10.18112/openneuro.ds003768.v1.0.0.

## Disclaimers

### Disclosures

Outside of the current work, Orfeu M. Buxton discloses that he received subcontract grants to Penn State from Proactive Life LLC (formerly Mobile Sleep Technologies) doing business as SleepScape (NSF/STTR #1622766, NIH/NIA SBIR R43-AG056250, R44-AG056250), received honoraria/travel support for lectures from Boston University, Boston College, Tufts School of Dental Medicine, New York University and Allstate, consulting fees from SleepNumber, and receives an honorarium for his role as the Editor in Chief of Sleep Health (sleephealthjournal.org).

The other authors declare no competing financial interests and have no conflict of interest to declare.

## References

Akeju, O., Loggia, M.L., Catana, C., Pavone, K.J., Vazquez, R., Rhee, J., Ramirez, V.C., Chonde, D.B., Izquierdo-Garcia, D., Arabasz, G., Hsu, S., Habeeb, K., Hooker, J.M., Napadow, V., Brown, E.N., Purdon, P.L., 2014. Disruption of thalamic functional connectivity is a neural correlate of dexmedetomidine-induced unconsciousness. Elife 3, e04499. https://doi.org/10.7554/eLife.04499

Bandettini, P.A., Wong, E.C., Hinks, R.S., Tikofsky, R.S., Hyde, J.S., 1992. Time course EPI of human brain function during task activation. Magn. Reson. Med. 25, 390–397. https://doi.org/https://doi.org/10.1002/mrm.1910250220

Baust, W., Bohnert, B., 1969. The regulation of heart rate during sleep. Exp. Brain Res. 7, 169–180. https://doi.org/10.1007/BF00235442

Berger, H., 1929. Über das Elektrenkephalogramm des Menschen. Arch. Psychiatr. Nervenkr. 87, 527–570. https://doi.org/10.1007/BF01797193

Birn, R.M., Diamond, J.B., Smith, M.A., Bandettini, P.A., 2006. Separating respiratory-variation-related fluctuations from neuronal-activity-related fluctuations in fMRI. Neuroimage 31, 1536–1548. https://doi.org/10.1016/j.neuroimage.2006.02.048

Birn, R.M., Murphy, K., Bandettini, P.A., 2008a. The effect of respiration variations on independent component analysis results of resting state functional connectivity. Hum. Brain Mapp. 29, 740–750. https://doi.org/https://doi.org/10.1002/hbm.20577

Birn, R.M., Murphy, K., Handwerker, D.A., Bandettini, P.A., 2009. fMRI in the presence of task-correlated breathing variations. Neuroimage 47, 1092–1104. https://doi.org/10.1016/j.neuroimage.2009.05.030

Birn, R.M., Smith, M.A., Jones, T.B., Bandettini, P.A., 2008b. The respiration response function: The temporal dynamics of fMRI signal fluctuations related to changes in respiration. Neuroimage 40, 644–654. https://doi.org/10.1016/j.neuroimage.2007.11.059

Biswal, B., FZ, Y., VM, H., JS, H., 1995. -Functional connectivity in the motor cortex of resting human brain using echo-planar MRI. Magn Reson Med 34, 537–541. https://doi.org/10.1002/mrm.1910340409

Boudreau, P., Yeh, W.-H., Dumont, G.A., Boivin, D.B., 2013. Circadian Variation of Heart Rate Variability Across Sleep Stages. Sleep 36, 1919–1928. https://doi.org/10.5665/sleep.3230

Britz, J., Van De Ville, D., Michel, C.M., 2010. BOLD correlates of EEG topography reveal rapid resting-state network dynamics. Neuroimage 52, 1162–1170. https://doi.org/https://doi.org/10.1016/j.neuroimage.2010.02.052

Brookes, M.J., Hale, J.R., Zumer, J.M., Stevenson, C.M., Francis, S.T., Barnes, G.R., Owen, J.P., Morris, P.G., Nagarajan, S.S., 2011a. Measuring functional connectivity using MEG: methodology and comparison with fcMRI. Neuroimage 56, 1082–1104. https://doi.org/10.1016/j.neuroimage.2011.02.054

Brookes, M.J., Woolrich, M., Luckhoo, H., Price, D., Hale, J.R., Stephenson, M.C., Barnes, G.R., Smith, S.M., Morris, P.G., 2011b. Investigating the electrophysiological basis of resting state networks using magnetoencephalography. Proc. Natl. Acad. Sci. 108, 16783–16788. https://doi.org/10.1073/pnas.1112685108

Buxton, R.B., Uluda□, K., Dubowitz, D.J., Liu, T.T., 2004. Modeling the hemodynamic response to brain activation. Neuroimage 23, 220–233. https://doi.org/10.1016/j.neuroimage.2004.07.013

Chang, C., Cunningham, J.P., Glover, G.H., 2009. Influence of heart rate on the BOLD signal: The cardiac response function. Neuroimage 44, 857–869. https://doi.org/10.1016/j.neuroimage.2008.09.029

Chang, C., Glover, G.H., 2009. Relationship between respiration, end-tidal CO2, and BOLD signals in resting-state fMRI. Neuroimage 47, 1381–1393. https://doi.org/10.1016/j.neuroimage.2009.04.048

Chang, C., Leopold, D.A., Scholvinck, M.L., Mandelkow, H., Picchioni, D., Liu, X., Ye, F.Q., Turchi, J.N., Duyn, J.H., 2016. Tracking brain arousal fluctuations with fMRI. Proc. Natl. Acad. Sci. U. S. A. 113, 4518–4523. https://doi.org/10.1073/pnas.1520613113

Chang, C., Metzger, C.D., Glover, G.H., Duyn, J.H., Heinze, H.J., Walter, M., 2013. Association between heart rate variability and fluctuations in resting-state functional connectivity. Neuroimage 68, 93–104. https://doi.org/10.1016/j.neuroimage.2012.11.038

Dampney, R.A.L., 2015. Central mechanisms regulating coordinated cardiovascular and respiratory function during stress and arousal. Am. J. Physiol. Integr. Comp. Physiol. 309, R429–R443. https://doi.org/10.1152/ajpregu.00051.2015

Das, A., Murphy, Kevin, Drew, P.J., Murphy, K, Pj, D., 2021. Rude mechanicals in brain haemodynamics□: non-neural actors that influence blood flow. Philos. Trans. R. Soc. B 376, 20190635.

de Munck, J.C., Gonçalves, S.I., Huijboom, L., Kuijer, J.P.A., Pouwels, P.J.W., Heethaar, R.M., Lopes da Silva, F.H., 2007. The hemodynamic response of the alpha rhythm: An EEG/fMRI study. Neuroimage 35, 1142–1151. https://doi.org/10.1016/j.neuroimage.2007.01.022

de Zambotti, M., Willoughby, A.R., Franzen, P.L., Clark, D.B., Baker, F.C., Colrain, I.M., 2016. K-Complexes: Interaction between the Central and Autonomic Nervous Systems during Sleep. Sleep 39, 1129–1137. https://doi.org/10.5665/sleep.5770

Douglas, N.J., White, D.P., Pickett, C.K., Weil, J. V, Clifford, W., 1982. Respiration during sleep in normal man 37, 840–844.

Drew, P.J., Mateo, C., Turner, K.L., Yu, X., Kleinfeld, D., 2020. Ultra-slow Oscillations in fMRI and Resting-State Connectivity□: Neuronal and Vascular Contributions and Technical Confounds. Neuron 107, 782–804. https://doi.org/10.1016/j.neuron.2020.07.020

Duyn, J.H., Ozbay, P.S., Chang, C., Picchioni, D., 2020. Physiological changes in sleep that affect fMRI inference. Curr. Opin. Behav. Sci. 33, 42–50. https://doi.org/10.1016/j.cobeha.2019.12.007

Falahpour, M., Chang, C., Wong, C.W., Liu, T.T., 2018. Template-based prediction of vigilance fluctuations in resting-state fMRI. Neuroimage 174, 317–327. https://doi.org/10.1016/j.neuroimage.2018.03.012

Feige, B., Scheffler, K., Esposito, F., Di Salle, F., Hennig, J., Seifritz, E., 2005. Cortical and subcortical correlates of electroencephalographic alpha rhythm modulation. J. Neurophysiol. 93, 2864–2872. https://doi.org/10.1152/jn.00721.2004

Fox, M.D., Raichle, M.E., 2007. Spontaneous fluctuations in brain activity observed with functional magnetic resonance imaging. Nat Rev Neurosci 8, 700–711. https://doi.org/nrn2201 [pii]\n10.1038/nrn2201

Fultz, N.E., Bonmassar, G., Setsompop, K., Stickgold, R.A., Rosen, B.R., Polimeni, J.R., Lewis, L.D., 2019. Coupled electrophysiological, hemodynamic, and cerebrospinal fluid oscillations in human sleep. Science (80-.). 366, 628–631. https://doi.org/10.1126/science.aax5440

Glasser, M.F., Coalson, T.S., Bijsterbosch, J.D., Harrison, S.J., Harms, M.P., Anticevic, A., Essen, D.C. Van, Smith, S.M., 2018. Using temporal ICA to selectively remove global noise while preserving global signal in functional MRI data. Neuroimage 181, 692–717. https://doi.org/10.1016/j.neuroimage.2018.04.076

Goldman, R., Stern, J., Jr, J.E., Cohen, M., 2002. Simultaneous EEG and fMRI of the alpha rhythm. Neuroreport 13, 2487–2492. https://doi.org/10.1097/01.wnr.0000047685.08940.d0.Simultaneous

Gu, Y., Han, F., Liu, X., 2019. Arousal Contributions to Resting-State fMRI Connectivity and Dynamics. Front. Neurosci. 13, 1190.

Gu, Y., Sainburg, L.E., Kuang, S., Han, F., Williams, J.W., Liu, Y., Zhang, N., Zhang, X., Leopold, D.A., Liu, X., 2021. Brain Activity Fluctuations Propagate as Waves Traversing the Cortical Hierarchy. Cereb. Cortex 1–20. https://doi.org/10.1093/cercor/bhab064

Han, F., Brown, G.L., Zhu, Y., Belkin-Rosen, A.E., Lewis, M.M., Du, G., Gu, Y., Eslinger, P.J., Mailman, R.B., Huang, X., Liu, X., 2021a. Decoupling of global brain activity and cerebrospinal fluid flow in Parkinson’s cognitive decline. Mov. Disord. https://doi.org/10.1101/2021.01.08.425953

Han, F., Chen, J., Belkin-Rosen, A., Gu, Y., Luo, L., Buxton, O.M., Liu, X., 2021b. Reduced coupling between cerebrospinal fluid flow and global brain activity is linked to Alzheimer disease–related pathology. PLOS Biol. 19, e3001233. https://doi.org/10.1371/journal.pbio.3001233

He, B.J., Snyder, A.Z., Zempel, J.M., Smyth, M.D., Raichle, M.E., 2008. Electrophysiological correlates of the brain{\textquoteright}s intrinsic large-scale functional architecture. Proc. Natl. Acad. Sci. 105, 16039–16044. https://doi.org/10.1073/pnas.0807010105

Iliff, J.J., Wang, M., Liao, Y., Plogg, B.A., Peng, W., Gundersen, G.A., Benveniste, H., Vates, G.E., Deane, R., Goldman, S.A., Nagelhus, E.A., Nedergaard, M., 2012. A Paravascular Pathway Facilitates CSF Flow Through the Brain Parenchyma and the Clearance of Interstitial Solutes, Including Amyloid β. Sci. Transl. Med. 4, 147ra111. https://doi.org/10.1126/scitranslmed.3003748

Iliff, J.J., Wang, M., Zeppenfeld, D.M., Venkataraman, A., Plog, B.A., Liao, Y., Deane, R., Nedergaard, M., 2013a. Cerebral Arterial Pulsation Drives Paravascular CSF-Interstitial Fluid Exchange in the Murine Brain. J. Neurosci. 33, 18190–18199. https://doi.org/10.1523/JNEUROSCI.1592-13.2013

Iliff, J.J., Wang, M., Zeppenfeld, D.M., Venkataraman, A., Plog, B.A., Liao, Y., Deane, R., Nedergaard, M., 2013b. Cerebral Arterial Pulsation Drives Paravascular CSF–Interstitial Fluid Exchange in the Murine Brain. J. Neurosci. 33, 18190–18199. https://doi.org/10.1523/JNEUROSCI.1592-13.2013

Keilholz, S.D., Pan, W.-J., Billings, J., Nezafati, M., Shakil, S., 2017. Noise and non-neuronal contributions to the BOLD signal: applications to and insights from animal studies. Neuroimage 154, 267–281. https://doi.org/https://doi.org/10.1016/j.neuroimage.2016.12.019

Klose, U., Strik, C., Kiefer, C., Grodd, W., 2000. Detection of a relation between respiration and CSF pulsation with an echoplanar technique. J. Magn. Reson. Imaging 11, 438–444. https://doi.org/https://doi.org/10.1002/(SICI)1522-2586(200004)11:4<438::AID-JMRI12>3.0.CO;2-O

Kwong, K.K., Belliveau, J.W., Chesler, D.A., Goldberg, I.E., Weisskoff, R.M., Poncelet, B.P., Kennedy, D.N., Hoppel, B.E., Cohen, M.S., Turner, R., 1992. Dynamic magnetic resonance imaging of human brain activity during primary sensory stimulation. Proc. Natl. Acad. Sci. U. S. A. 89, 5675–9. https://doi.org/10.1073/pnas.89.12.5675

Liu, X., De Zwart, J.A., Schölvinck, M.L., Chang, C., Ye, F.Q., Leopold, D.A., Duyn, J.H., 2018. Subcortical evidence for a contribution of arousal to fMRI studies of brain activity. Nat. Commun. 9, 1–10. https://doi.org/10.1038/s41467-017-02815-3

Liu, X., Leopold, D.A., Yang, Y., 2021. Single neuron firing cascades underlie global spontaneous brain events. BioRxiv 1–33.

Liu, X., Yanagawa, T., Leopold, D.A., Chang, C., Ishida, H., Fujii, N., Duyn, J.H., 2015. Arousal transitions in sleep, wakefulness, and anesthesia are characterized by an orderly sequence of cortical events. Neuroimage 116, 222–231. https://doi.org/10.1016/j.neuroimage.2015.04.003

Liu, Z., de Zwart, J.A., Yao, B., van Gelderen, P., Kuo, L.W., Duyn, J.H., 2012. Finding thalamic BOLD correlates to posterior alpha EEG. Neuroimage 63, 1060–1069. https://doi.org/10.1016/j.neuroimage.2012.08.025

Majeed, W., Magnuson, M., Hasenkamp, W., Schwarb, H., Schumacher, E.H., Barsalou, L., Keilholz, S.D., 2011. Spatiotemporal dynamics of low frequency BOLD fluctuations in rats and humans. Neuroimage 54, 1140–1150. https://doi.org/10.1016/j.neuroimage.2010.08.030

Makeig, S., Inlow, M., 1993. Lapse in alertness: coherence of fluctuations in performance and EEG spectrum. Electroencephalogr. Clin. Neurophysiol. 86, 23–35. https://doi.org/https://doi.org/10.1016/0013-4694(93)90064-3

Mantini, D., Perrucci, M.G., Gratta, C. Del, Romani, G.L., Corbetta, M., 2007. Electrophysiological signatures of resting state networks in the human brain. Proc Natl. Acad. Sci. USA 104, 13170–13175. https://doi.org/10.1073/pnas.0700668104

Mitra, A., Snyder, A.Z., Blazey, T., Marcus, E., 2015. Correction for Mitra et al., Lag threads organize the brain’s intrinsic activity. Proc. Natl. Acad. Sci. 112, E7307–E7307. https://doi.org/10.1073/pnas.1523893113

Moosmann, M., Ritter, P., Krastel, I., Brink, A., Thees, S., Blankenburg, F., Taskin, B., Obrig, H., Villringer, A., 2003. Correlates of alpha rhythm in functional magnetic resonance imaging and near infrared spectroscopy. Neuroimage 20, 145–158. https://doi.org/10.1016/S1053-8119(03)00344-6

Musso, F., Brinkmeyer, J., Mobascher, A., Warbrick, T., Winterer, G., 2010. Spontaneous brain activity and EEG microstates. A novel EEG/fMRI analysis approach to explore resting-state networks. Neuroimage 52, 1149–1161. https://doi.org/https://doi.org/10.1016/j.neuroimage.2010.01.093

Ogawa, S., Tank, D.W., Menon, R., Ellermann, J.M., Kim, S.G., Merkle, H., Ugurbil, K., 1992. Intrinsic signal changes accompanying sensory stimulation: functional brain mapping with magnetic resonance imaging. Proc. Natl. Acad. Sci. 89, 5951–5955. https://doi.org/10.1073/pnas.89.13.5951

Özbay, P.S., Chang, C., Picchioni, D., Mandelkow, H., Chappel-Farley, M.G., van Gelderen, P., de Zwart, J.A., Duyn, J., 2019. Sympathetic activity contributes to the fMRI signal. Commun. Biol. 2, 1–9. https://doi.org/10.1038/s42003-019-0659-0

Özbay, P.S., Chang, C., Picchioni, D., Mandelkow, H., Moehlman, T.M., Chappel-Farley, M.G., van Gelderen, P., de Zwart, J.A., Duyn, J.H., 2018. Contribution of systemic vascular effects to fMRI activity in white matter. Neuroimage 176, 541–549. https://doi.org/10.1016/j.neuroimage.2018.04.045

Picchioni, D., Özbay, P.S., Mandelkow, H., de Zwart, J.A., Wang, Y., van Gelderen, P., Duyn, J.H., 2021. Autonomic arousals contribute to brain fluid pulsations during sleep. bioRxiv 2021.05.04.442672. https://doi.org/10.1101/2021.05.04.442672

Power, J.D., Barnes, K.A., Snyder, A.Z., Schlaggar, B.L., Petersen, S.E., 2012. Spurious but systematic correlations in functional connectivity MRI networks arise from subject motion. Neuroimage 59, 2142–2154. https://doi.org/10.1016/j.neuroimage.2011.10.018

Power, J.D., Schlaggar, B.L., Petersen, S.E., 2015. Recent progress and outstanding issues in motion correction in resting state fMRI. Neuroimage 105, 536–551. https://doi.org/10.1016/j.neuroimage.2014.10.044

Putilov, A.A., Donskaya, O.G., 2014. Alpha attenuation soon after closing the eyes as an objective indicator of sleepiness. Clin. Exp. Pharmacol. Physiol. 41, 956–964. https://doi.org/https://doi.org/10.1111/1440-1681.12311

Raichle, M.E., 2015. The Brain’ s Default Mode Network. Annu. Rev. Neurosci. 38, 433–447. https://doi.org/10.1146/annurev-neuro-071013-014030

Raichle, M.E., Macleod, A.M., Snyder, A.Z., Powers, W.J., Gusnard, D.A., Shulman, G.L., 2001. A default mode of brain function. Proc Natl. Acad. Sci. USA 98, 676–682.

Sadaghiani, S., D’Esposito, M., 2015. Functional characterization of the cingulo-opercular network in the maintenance of tonic alertness. Cereb. Cortex 25, 2763–2773. https://doi.org/10.1093/cercor/bhu072

Schley, D., Carare-Nnadi, R., Please, C.P., Perry, V.H., Weller, R.O., 2006. Mechanisms to explain the reverse perivascular transport of solutes out of the brain. J. Theor. Biol. 238, 962–974. https://doi.org/https://doi.org/10.1016/j.jtbi.2005.07.005

Scholvinck, M.L., Maier, A., Ye, F.Q., Duyn, J.H., Leopold, D.A., 2010. Neural basis of global resting-state fMRI activity. Proc. Natl. Acad. Sci. 107, 10238–10243. https://doi.org/10.1073/pnas.0913110107

Seeley, W.W., Menon, V., Schatzberg, A.F., Keller, J., Glover, G.H., Kenna, H., Reiss, A.L., Greicius, M.D., 2007. Dissociable intrinsic connectivity networks for salience processing and executive control. J. Neurosci. 27, 2349–2356. https://doi.org/10.1523/JNEUROSCI.5587-06.2007

Shmueli, K., van Gelderen, P., de Zwart, J.A., Horovitz, S.G., Fukunaga, M., Jansma, J.M., Duyn, J.H., 2007. Low-frequency fluctuations in the cardiac rate as a source of variance in the resting-state fMRI BOLD signal. Neuroimage 38, 306–320. https://doi.org/10.1016/j.neuroimage.2007.07.037

Silvani, A., Calandra-Buonaura, G., Benarroch, E.E., Dampney, R.A.L., Cortelli, P., 2015. Bidirectional interactions between the baroreceptor reflex and arousal: an update. Sleep Med. 16, 210–216. https://doi.org/https://doi.org/10.1016/j.sleep.2014.10.011

Snyder, F., Hobson, J.A., Morrison, D.F., Goldfrank, F., 1964. Changes in respiration, heart rate, and systolic blood pressure in human sleep. J. Appl. Physiol. 19, 417–422. https://doi.org/10.1152/jappl.1964.19.3.417

Soon, C.S., Vinogradova, K., Ong, J.L., Calhoun, V.D., Liu, T., Zhou, J.H., Ng, K.K., Chee, M.W.L., 2021. Respiratory, cardiac, EEG, BOLD signals and functional connectivity over multiple microsleep episodes. Neuroimage 237, 118129. https://doi.org/https://doi.org/10.1016/j.neuroimage.2021.118129

Stoodley, M.A., Brown, S.A., Brown, C.J., Jones, N.R., 1997. Arterial pulsation—dependent perivascular cerebrospinal fluid flow into the central canal in the sheep spinal cord. J. Neurosurg. 86, 686–693. https://doi.org/10.3171/jns.1997.86.4.0686

Tarasoff-Conway, J.M., Carare, R.O., Osorio, R.S., Glodzik, L., Butler, T., Fieremans, E., Axel, L., Rusinek, H., Nicholson, C., Zlokovic, B. V, Frangione, B., Blennow, K., Ménard, J., Zetterberg, H., Wisniewski, T., de Leon, M.J., 2015. Clearance systems in the brain-implications for Alzheimer disease. Nat. Rev. Neurol. 11, 457–470. https://doi.org/10.1038/nrneurol.2015.119

Tong, Y., Hocke, L.M., Frederick, B.B., Chen, J., 2019a. Low Frequency Systemic Hemodynamic “Noise” in Resting State BOLD fMRI□: Characteristics, Causes, Implications, Mitigation Strategies, and Applications. Front. Neurosci. 13. https://doi.org/10.3389/fnins.2019.00787

Tong, Y., Hocke, L.M., Licata, S.C., de B. Frederick, B., 2012. Low-frequency oscillations measured in the periphery with near-infrared spectroscopy are strongly correlated with blood oxygen level-dependent functional magnetic resonance imaging signals. J. Biomed. Opt. 17, 106004. https://doi.org/10.1117/1.jbo.17.10.106004

Tong, Y., Yao, J. (Fiona), Chen, J.J., Frederick, B. de B., 2019b. The resting-state fMRI arterial signal predicts differential blood transit time through the brain. J. Cereb. Blood Flow Metab. 39, 1148–1160. https://doi.org/10.1177/0271678X17753329

van Veluw, S.J., Hou, S.S., Calvo-Rodriguez, M., Arbel-Ornath, M., Snyder, A.C., Frosch, M.P., Greenberg, S.M., Bacskai, B.J., 2020. Vasomotion as a Driving Force for Paravascular Clearance in the Awake Mouse Brain. Neuron 105, 549–561.e5. https://doi.org/10.1016/j.neuron.2019.10.033

Xie, L., Kang, H., Xu, Q., Chen, M.J., Liao, Y., Thiyagarajan, M., Donnell, J.O., Christensen, D.J., Nicholson, C., Iliff, J.J., Takano, T., Deane, R., Nedergaard, M., 2013. Sleep Drives Metabolite Clearance from the Adult Brain. Science (80-.). 342, 373–378.

Yamada, S., Miyazaki, M., Yamashita, Y., Ouyang, C., Yui, M., Nakahashi, M., Shimizu, S., Aoki, I., Morohoshi, Y., McComb, J.G., 2013. Influence of respiration on cerebrospinal fluid movement using magnetic resonance spin labeling. Fluids Barriers CNS 10, 36. https://doi.org/10.1186/2045-8118-10-36

Yuan, H., Zotev, V., Phillips, R., Bodurka, J., 2013. Correlated slow fluctuations in respiration, EEG, and BOLD fMRI. Neuroimage 79, 81–93. https://doi.org/10.1016/j.neuroimage.2013.04.068

Zhang, D., Raichle, M.E., 2010. Disease and the brain’s dark energy. Nat. Rev. Neurol. 6, 15–28. https://doi.org/10.1038/nrneurol.2009.198

