## Supplementary material for "An orderly sequence of autonomic and neural events at transient arousal changes": Fig. S1, Fig. S2, Fig. S3, Fig. S4,Fig. S5, Fig. S6

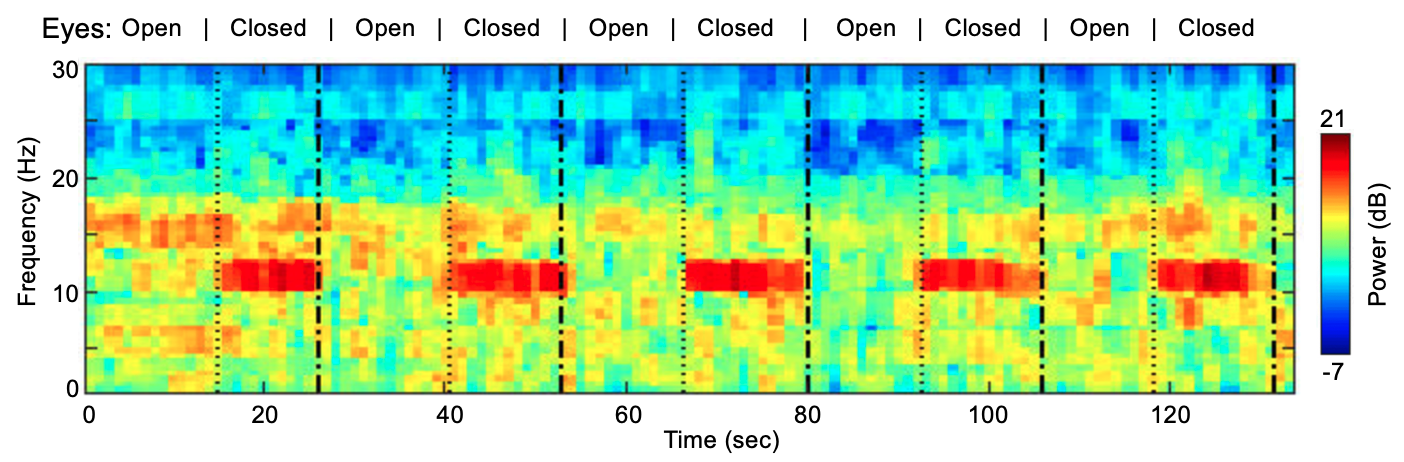

Fig. S1. A representative example of alpha power modulation under eyes-closed and eyes-open conditions from a subject. The spectrogram was averaged across the three occipital channels (O1, O2 and Oz). The dotted lines represent the moment when the subject pressed the button and started to close eyes and the dot-dashed liens represent the moment when the subject pressed the button and started to open eyes.

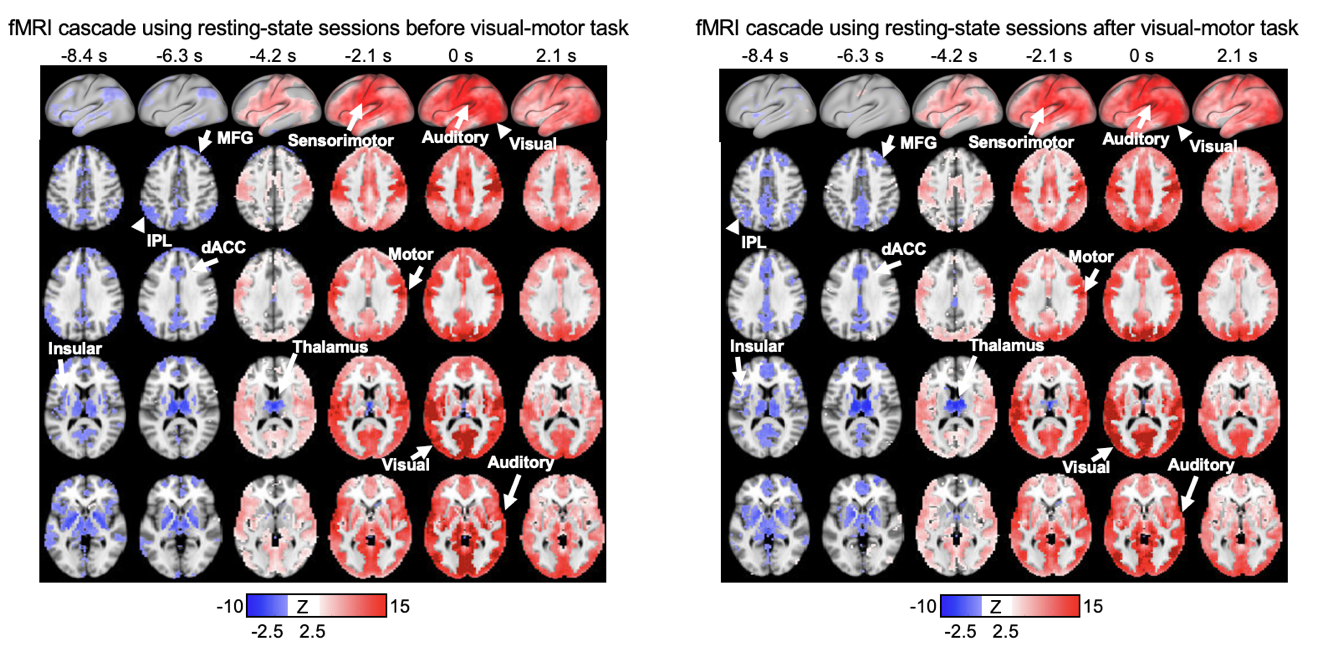

Fig. S2. The fMRI cascade calculated using resting-state sessions before (left) or after (right) the visual-motor adaptation task, respectively. The spatiotemporal pattern was converted to z-scored map and thresholded at an FDR-corrected q value of 0.01 for both maps. dACC, dorsal anterior cingulate cortex. IPL, inferior parietal lobe. MFG, medial frontal gyrus MFG. PCC, posterior cingulate cortex.

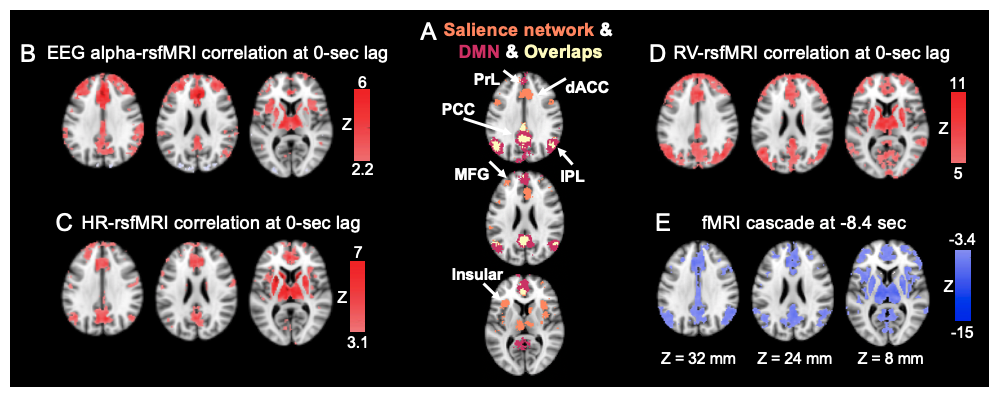

Fig. S3. The fMRI cascade and the rsfMRI correlations with various signals showed a spatial pattern resembling the salience and default mode networks. (A) The masks of the salience network or the default mode network were acquired from a Neurosynth meta-analysis (Yarkoni et al., 2011), by entering the term “salience network” or “default mode”. (B-D) The rsfMRI correlations with various signals at a time lag of 0 sec. (E) The fMRI cascade at -8.4 sec. PrL, prelimbic cortex.

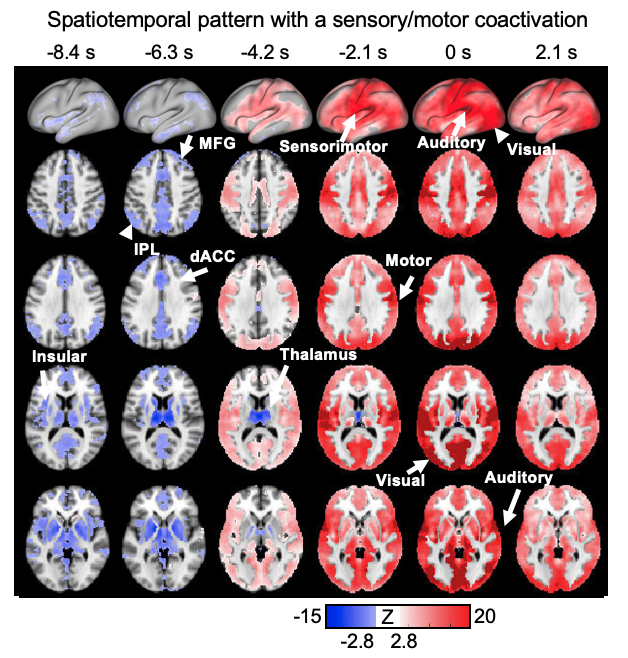

Fig. S4. The spatiotemporal pattern with sensory/motor co-activation pattern was calculated by averaged fMRI time segments around a sensory/motor co-activation pattern (N=329). The spatiotemporal pattern was converted to z-scored map and thresholded at an FDR-corrected q value of 0.005.

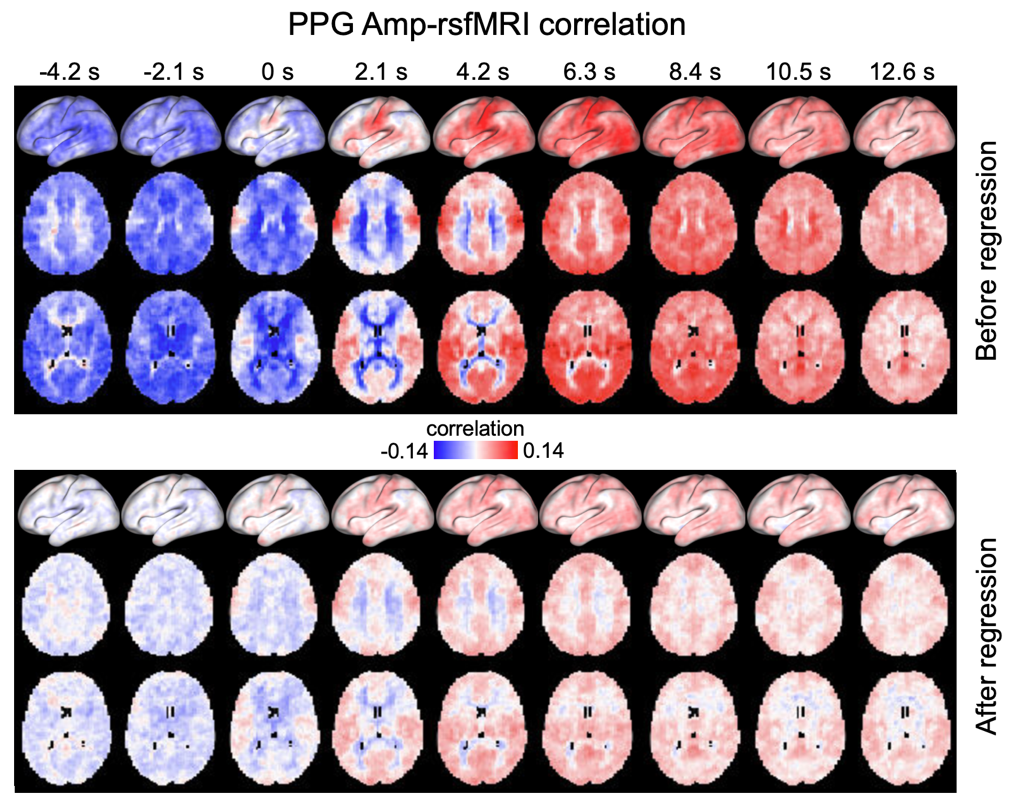

Fig. S5. The lag-dependent rsfMRI correlations with the PPG amplitude before (top) and after (bottom) regressing out the fMRI cascade.

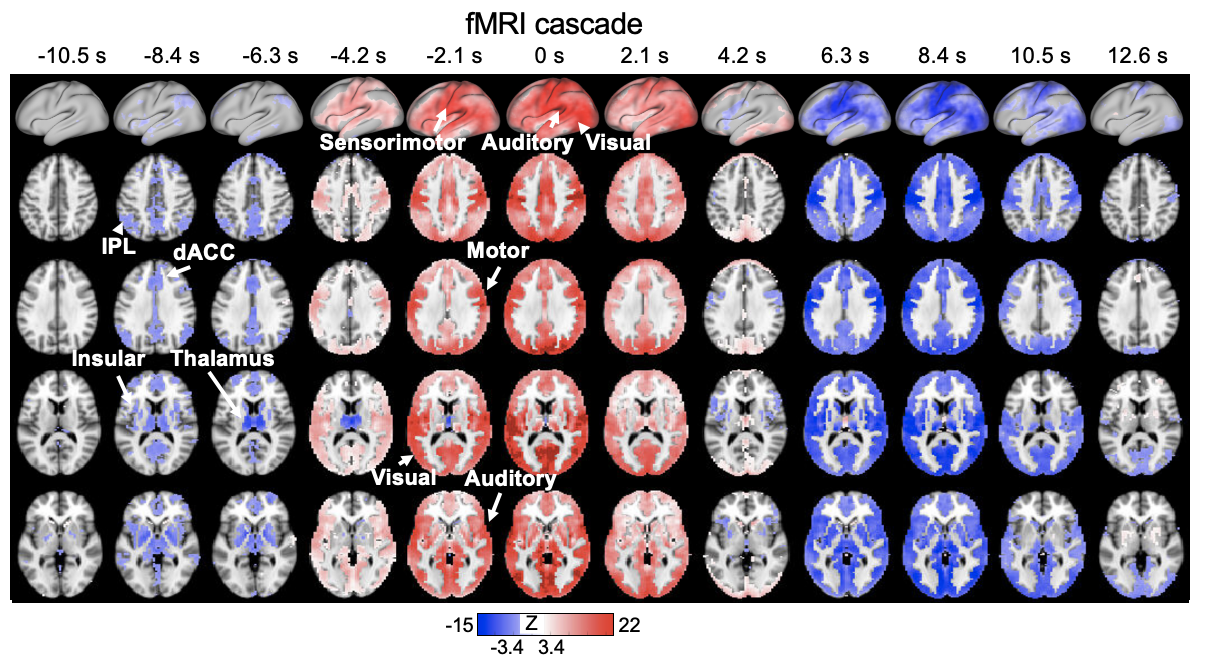

Fig. S6. The longer fMRI cascade with more time points included compared to that in Fig. 2A.
